# Wireless Sensor Network: New Concept of Spatial-Temporal Monitoring Plant–Environment Interactions

**DOI:** 10.1101/2025.09.04.674105

**Authors:** Idan Ifrach, Itamar Shenhar, Nir Averbuch, Bnaya Hami, Assaf Hurwitz, Menachem Moshelion

## Abstract

We present a low-cost, standards-based wireless sensor network (WSN) for continuous, canopy-integrated monitoring of plant–environment interactions. Each plant carries in-canopy microclimate sensors (temperature, relative humidity, illuminance) paired with nearby ambient references, yielding real-time canopy-ambient differentials. The system is easy to install: at planting or sowing, sensors are fixed at positions that will lie within the developing canopy, and a separate ambient reference area is designated and kept free of vegetation. As plants grow, they envelop the sensors, thereby capturing growth dynamics over time. The sensors accuracy was validated against a commercial weather station and portable system that measures gas exchange, temperature and light (LI-COR 6800/6400), and the system’s ability to resolve plant physiological activity was confirmed using the PlantArray functional phenotyping platform with independent whole-plant transpiration and biomass references. Under controlled growth-room conditions and across two contrasting Cannabis cultivars, daily transpiration strongly predicted biomass gain (R² > 0.9). Microclimate signals mirrored physiology: midday canopy air was cooler by 4–7 °C, more humid by 18–25 % RH, and increasingly shaded as biomass accumulated, with temperature, RH, and light attenuation showing saturating logarithmic relationships with growth. The network operated for months unattended with low packet loss and predictable power use. It provides 4D (x–y–z–time) coverage, where x and y denote horizontal location, z the vertical position within the canopy, and time the dynamics, enabling resolution of where changes occur and how they evolve, and supplying high-frequency labeled data. This system complements, rather than replaces, precision instruments and high-end phenotyping platforms, providing a scalable layer for continuous tracking across wide areas. We outline practical constraints and next steps toward field pilots, modest energy harvesting, expanded sensor suites, and integration with machine learning for predictive crop management.

## Introduction

Conventional farming methods typically apply uniform amounts of fertilizers, pesticides, and irrigation across entire fields, relying on limited sampling. This “blanket” approach, often based on the most stressed (e.g. nutrient-deficient areas), results in over-application elsewhere. Such inefficiency disregards the inherent small-scale variability in soil and microclimate conditions (Pierce & Nowak, 1999). In response, there is growing interest in advanced technologies and computational tools that enable early detection of biotic and abiotic plant stress and site-specific input application, which aims to optimize productivity by delivering resources precisely where and when they are needed. This paradigm lies at the core of Precision Agriculture (PA; Zhang et al., 2002). A central component of PA is remote sensing, which uses cameras and other sensors to collect data on soils, crops, and the atmosphere. Modern imaging platforms, spanning multispectral and hyperspectral remote sensing from satellites and aircraft to low-altitude, together with proximal RGB cameras and LiDAR, provide high-resolution spatial monitoring of canopy status, structural traits, and three-dimensional field reconstructions (Chebrolu et al., 2021a; Li et al., 2014). Yet, despite their breadth of coverage and rich spatial detail, these systems typically acquire data at discrete time points dictated by orbital revisit, flight logistics, or scanning schedules, yielding snapshots rather than continuous records, which limits the timely detection of rapid physiological dynamics during the growing season (Chebrolu et al., 2021b; Li et al., 2014). While spatiotemporal modeling can partially bridge intervals between acquisitions, the underlying measurement cadence remains discrete (Pérez-Valencia et al., 2022).

Capturing rapid physiological dynamics would require impractically frequent acquisitions, underscoring the need for complementary systems that can deliver continuous, canopy-integrated measurements.

### Physiological Basis for High-Frequency Monitoring

Plants can undergo rapid physiological changes, highlighting the risk of missing critical events when measurements are infrequent. A prime example is stomatal behavior, as stomatal responses to environmental fluctuations are among the earliest indicators (minutes) of a plant’s status in balancing productivity and survival (Lupo & Moshelion, 2024). Plant productivity depends on CO₂ uptake through open stomata, enabling photosynthesis using sunlight and soil water.

However, this gas exchange comes at the cost of water loss, as most water absorbed by the roots exits through transpiration via the same stomata. Transpiration is beneficial while soil moisture is sufficient, as it facilitates nutrient along with water transport and leaf cooling. Under water-limited conditions, plants rapidly reduce stomatal opening, thereby limiting both carbon fixation and transpiration, shifting from a productivity-oriented mode to a survival state (Moshelion et al., 2024). Additionally, stomatal closure also occurs in response to biotic stress. Studies have shown that stomata can close following pathogen attack days before visible symptoms appear (Fortin & Friedman, 2025). Therefore, continuous monitoring of stomatal behavior, or related traits such as transpiration, can provide early warning of diverse stress conditions well before visual symptoms like wilting or color change become apparent.

Stomatal and other physiological adjustments occur on short timescales. Studies have shown that stomatal conductance can respond to environmental fluctuations, such as changes in light intensity, within minutes (Matthews et al., 2018; Tanaka et al., 2019). This rapid responsiveness is essential for continuous optimization of plant performance and has direct implications for growth and yield (Shenhar et al., 2025). The momentary reactivity of stomata is driven by complex sensory and signaling networks within guard cells. For example, changes in light are detected by photoreceptors (Breen et al., 2023), temperature by thermo-sensors (Kerbler and Wigge, 2023), atmospheric CO₂ by CO₂ receptors (Tian et al., 2015), and drought or salinity by phytohormonal and electrical signals (Baxter et al., 2014; Waadt et al., 2022). Nutrient availability also modulates signaling pathways, including those involving reactive oxygen species (ROS) and turgor regulation, which influence stomatal behavior (Schachtman and Shin, 2007). In particular, nitrogen deficiency reduces daily transpiration even under irrigation (Weksler et al., 2021a), while potassium deficiency disrupts stomatal regulation and may increase water loss under stress (Weksler et al., 2021b). These real-time stomatal responses to environmental signals lead to dynamic shifts in gas exchange and transitions between growth and survival modes. Therefore, continuous monitoring of key physiological traits, paired with real-time analysis, can offer early diagnostics of plant stress well before visual symptoms appear. Because these responses unfold on minute-to-hour intervals, effective diagnosis requires high-frequency, continuous, canopy-integrated sensing, which brings into focus the spatiotemporal trade-off in current field methods.

### Trade-Off Between Manual and Remote Sensing in Spatial-Temporal Resolution

Current plant monitoring methods face a trade-off between measurement accuracy and coverage in both time and space. This trade-off, often referred to as spatiotemporal (spatial-temporal) resolution, reflects the balance between how precisely a sensor can detect variation across an area and how frequently it can measure changes at the same location. Manual physiological measurements, such as leaf gas exchange or water potential, provide accurate snapshots of plant water status. However, these measurements are infrequent, labor-intensive, and limited in throughput, typically representing only one leaf or plant per reading. As such, they are not suitable for high-frequency monitoring across large populations. By contrast, camera-based remote/proximal sensing scales to whole experiments and enables rapid acquisitions, yet measurements occur at discrete time points that yield snapshots rather than continuous records; analyses therefore often rely on interpolation between acquisitions (Chebrolu et al., 2021b). This gap at minute-to-hour time scales constrains the early detection of stress at population scale. To overcome this spatiotemporal trade-off and capture minute-scale plant responses in situ, we implemented an “always-on”, canopy-integrated wireless sensor network.

### Wireless Sensor Network for Continuous Plant–Environment Monitoring

Wireless sensor networks (WSNs) enable the deployment of distributed sensors to monitor environmental variability at high spatial and temporal resolution, with high precision and reasonable costs (see Materials and Methods). These telemetric systems consist of stationary sensor nodes, densely arranged in a flexible grid, that autonomously measure and transmit environmental data in real time (Akyildiz et al., 2002; Bormann et al., 2014). WSNs, enable in situ observation at plant and/or ground level, offering self-organizing and tolerant architectures that address spatiotemporal sensing challenges beyond the reach of conventional remote platforms (Aqeel-Ur-Rehman et al., 2014; Glasgow et al., 2004; Romanov et al., 2016). For example, WSNs have been deployed to monitor greenhouse microclimate for disease control in tomato crops (Mancuso and Bustaffa, 2006) and to operate solar-powered irrigation in outdoor farms in Malawi (Mafuta et al., 2013). Broader applications include habitat surveillance and open-field monitoring in vineyards, paprika systems, and large-scale agricultural settings (Beckwith et al., 2004; Hwang et al., 2010; Lloret et al., 2011; Mainwaring et al., 2002; Romanov et al., 2016; Zhao et al., 2011). However, despite their well-established role in environmental monitoring, WSNs have rarely been deployed to capture real-time physiological feedback from plants to their ambient conditions. While these systems efficiently monitor abiotic factors, their application to internal plant processes, such as transpiration and growth rate, remains limited, reflecting the current technological challenges associated with continuous, in-field deployment (Großkinsky et al., 2015; Kumar et al., 2022). Additionally, in field conditions, sensors must operate continuously and autonomously, often for an entire growing season, preferably without battery replacement. This requirement compels a balance between energy-saving strategies and high-frequency, reliable data acquisition. Frequent data transmission is essential for capturing minute-scale physiological responses to environmental fluctuations, yet high transmission rates increase energy consumption and shorten battery life. Conversely, low-frequency transmission risks missing critical physiological events. Off-the-shelf Bluetooth options illustrate the trade-offs. Classic Bluetooth star topologies support up to ∼7 active nodes per router. Bluetooth Mesh, introduced in 2017, theoretically scales to tens of thousands of nodes, but in dense deployments flooding-based relays and interference degrade throughput/latency and overall performance (Hortelano et al., 2017b; Iii et al., 2016; Nikodem et al., 2020; Nikodem & Bawiec, 2019). Energetically, high-frequency transmissions and larger payloads raise consumption, whereas very small packets cap throughput (Hortelano et al., 2017b, 2017a; Kim et al., 2015). In addition, Bluetooth’s service model can limit flexibility: the Bluetooth SIG defines standardized service identifiers (UUIDs, unique 128-bit IDs for services/characteristics). Vendor stacks often restrict custom UUIDs and proprietary commands, pushing reliance on standard profiles and closed-source implementations; in practice this constrains scalability beyond ∼100 nodes and challenges multi-month telemetry (Hortelano et al., 2017a; Spörk et al., 2017). By contrast, to overcome these limitations we adopted a standards-based, low-power IPv6 approach (IETF 6TiSCH; see Methods), which schedules brief transmit/receive slots and hops channels to reduce interference while keeping radios asleep between slots, enabling long-lived, high-integrity telemetry at scale. In practice, the IETF 6TiSCH family of protocols has been validated in large testbeds and is known to provide stable, scalable, and energy-efficient communication (Duquennoy et al., 2017; Thubert, 2021; Vilajosana et al., 2020). At a high level, the network coordinates when nodes transmit and hops among radio channels, which reduces interference and saves energy, advantages that are important for plant–environment monitoring at scale. In this study, we introduce a novel concept for a plant–environment sensor network designed to provide high spatiotemporal resolution monitoring. This approach is essential for accurately capturing dynamic plant water-balance responses and optimizing crop management under variable conditions. Additionally, we propose a new approach that enables continuous, detailed monitoring of plant–environment interactions, tracking the subtle environmental changes induced by plants in their immediate surroundings. Specifically, we suggest than continuedly monitoring the microclimate within the plant canopy alongside a nearby reference point that remains unaffected by the plant. This paired sensing strategy may provide a dynamic view of plant physiological activity through its impact on local conditions. This framework forms the basis for our central hypothesis, linking plant developmental and physiological processes to quantifiable microclimatic signatures.

### Hypothesis

We hypothesize that a plant’s physiological responses and developmental rate can be quantitatively inferred by measuring the microclimate differences between the plant canopy and a nearby reference point. By continuously tracking these in-canopy versus ambient deltas, we expect to obtain real-time indicators of plant activity, stress status, and developmental progression relative to other plants. In this context, larger microclimatic deltas are interpreted as indicators of higher physiological activity.

In the short term (minutes to hours), increased transpiration is expected to cool the air within the canopy and elevate relative humidity, creating a microclimate that is cooler and more humid than the surrounding ambient air. These short-term microclimate changes serve as real-time proxies for stomatal conductance and photosynthetic activity.

We propose that this high-frequency, in situ approach can serve as a non-invasive surrogate for traditional physiological measurements, capturing plant responses across multiple timescales.

## Materials and Methods

### Study Site and Plant Material

*Cannabis* sativa L. (Type I chemotype) was selected as the model species, using two commercial cultivars, ‘Odem’ and ‘MVA’, chosen for their contrasting canopy architectures. ‘Odem’ exhibits a tall, sativa-like structure, while ‘MVA’ has a more compact, indica-like morphology (Figure 1F and 1G). To minimize genetic variability, all experimental plants were propagated from single mother plants for each cultivar. Rooted cuttings, three weeks old, were transplanted into 7-L pots filled with a peat-based substrate (”540”, Kekkila, Finland) and grown under uniform conditions. Nutrients were delivered through automated fertigation using a customized solution (Super Shefer, ICL Fertilizers, Israel), supplemented with calcium and magnesium, and applied via proportional dosing pumps (DDA 5-00, Dosatron, France) at each irrigation. Experiments were conducted in a controlled-environment growth room (∼25 m²) at the R.H. Smith Faculty of Agriculture, Hebrew University of Jerusalem (Rehovot campus, Israel). Wireless sensor nodes were installed on a subset of the plants included in the broader PlantArray phenotyping study described by Shenhar et al. (2025).

**Figure 1.**
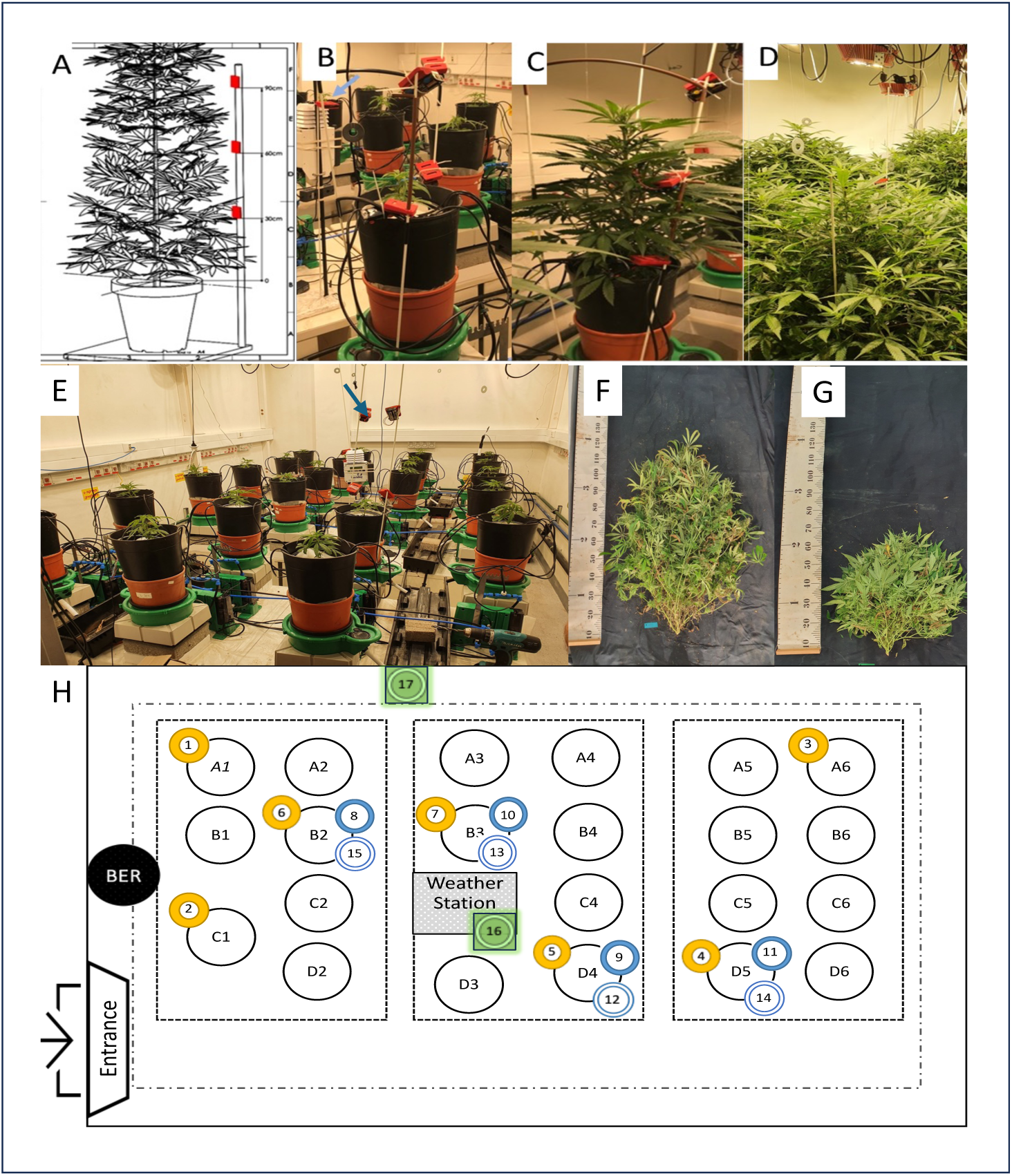
Spatial arrangement and installation of sensor platforms (SPs) in the experimental growth room. (A) Schematic illustration showing the vertical placement of three SP units per plant at 30, 60, and 90 cm above the pot base. (B) Sensor configuration on the planting day: the lower SP is partially visible, while mid and upper SPs are fully exposed above the foliage. (C) Twelve days after planting, lower SPs are mostly concealed by expanding foliage. (D) By 30 days post-planting, both lower and mid-canopy SPs are completely covered, while upper SPs remain exposed above the canopy (see also the individually adjustable lamp system installed for each plant). (E) Overview of the controlled-environment growth room on planting day, showing 22 plants installed on individual load-cell lysimeters (PlantArray 3.0). The WatchDog weather station (blue arrow) is positioned at the room center for continuous logging of ambient atmospheric conditions. (F–G) Post-harvest comparison of canopy structure in the two cultivars used: ‘Odem’ (F, tall sativa-type) and ‘MVA’ (G, compact indica-type). (H) Spatial layout of the wireless sensor network (WSN), including 17 SPs. Fifteen SPs were installed on seven selected plants (A01, A06, B02, B03, C01, D04, D05), each with a 30 cm SP (yellow circles); four plants had additional SPs at 60 cm (filled blue circles), and four also at 90 cm (hollow blue circles). Two additional ambient reference SPs were installed: one near the WatchDog weather station (#16) and another near the wall (#17).

Plants were cultivated from the vegetative stage through harvest in the same growth room. Environmental conditions, including light intensity and photoperiod, were regulated to simulate commercial cultivation protocols. Illumination was provided by overhead LED fixtures (Cree CXB3590, 3000 K + 4000 K spectrum), delivering a mean photosynthetic photon flux density (PPFD) of ∼1200 µmol m⁻² s⁻¹ at the canopy top. A long-day photoperiod (16 h light / 8 h dark) was used to promote vegetative growth, to simulate a daily sun spectrum variation the first two hours each day start with 4000K spectrum, procced with both spectrum and finished with 2 hours of 3000K spectrum (Figure 2A). At the beginning of the growth period, the fixtures were raised every 14–20 days, and as plant growth accelerated, the adjustment frequency increased to every few days. The adjustment system allowed individual height control for each plant, ensuring a constant light intensity relative to the canopy height of each specimen (see Figure 1D).

**Figure 2.**
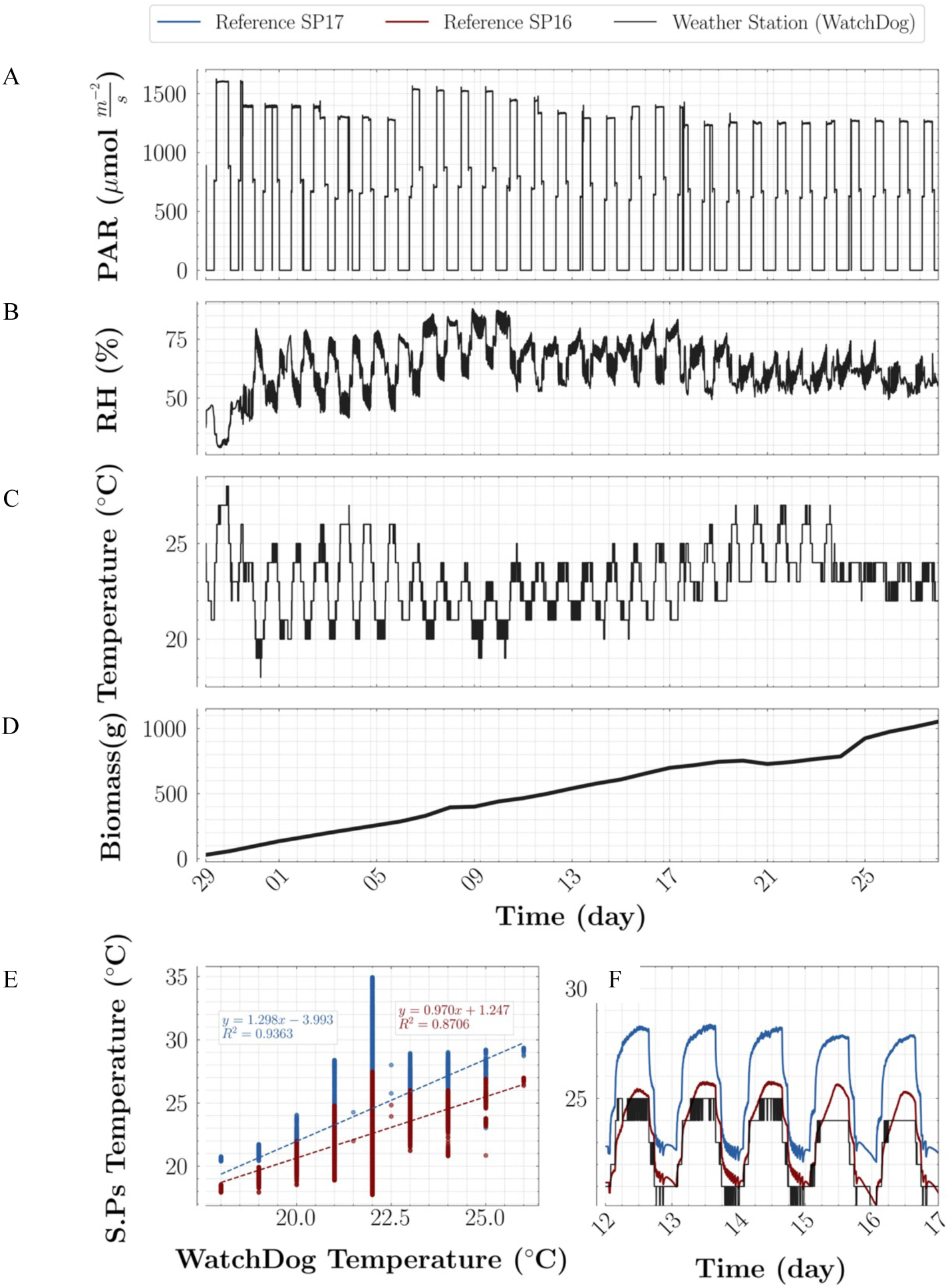
Continuous monitoring of environmental conditions and plant growth in the growth room. (A) Photosynthetically active radiation (PAR, μmol m⁻² s⁻¹), (B) relative humidity (RH, %), and (C) air temperature (°C) measured by the WatchDog weather station. (D) Plant biomass (g) of a representative plant, derived from net pot weight using the PlantArray system. (E) Correlation between temperature measurements from the WatchDog station and two reference sensor platforms (SP16 and SP17) over six consecutive days, demonstrating strong agreement (R² values shown). (F) Representative daily temperature patterns for the same period, showing consistent diurnal dynamics across these sensors.

Air temperature and relative humidity were controlled using active climate regulation, as well as four wall-mounted fans to minimize spatial heterogeneity within the room. Fans were oriented to promote uniform cross-canopy airflow between plants, thereby improving air mixing and achieving a spatially homogeneous microclimate. Environmental conditions were maintained similar to those reported in Shenhar et al. (2025), with temperature ranging from 20–29 °C (night–day, respectively) and relative humidity between 40–75% (day–night, respectively). A commercial weather station (WatchDog 2745, Spectrum Technologies, USA) was placed at the center of the room to continuously log ambient air temperature, relative humidity, and PPFD as the reference conditions (see Figure 1E; Figure 2 A-C). All plants were monitored using a functional phenotyping platform (PlantArray 3.0, Plant-DiTech, Israel), which enables high-throughput, real-time tracking of whole-plant water use efficiency, transpiration, and biomass accumulation. Each potted plant was placed on an individual load-cell lysimeter to allow precise gravimetric measurements. Data collection and calculations followed standard protocols described by Halperin et al. (2017), Dalal et al. (2020), and Shenhar et al. (2025).

### Whole Plant Physiological Measurements Using the Functional Phenotyping Platform

Whole-plant physiological traits were measured using a functional phenotyping system (PlantArray; Plant-DiTech Ltd., Yavne, Israel), which integrates high-precision, temperature-compensated load cells operating as weighing lysimeters. Each unit was connected to a controller that continuously collected pot-weight data and regulated irrigation according to plant transpiration. Data were logged every three minutes and accessed via web-based software.

Whole-plant physiological parameters were calculated following established protocols (Dalal et al., 2020; Halperin et al., 2017). **Daily Transpiration (DT):** the total daily water flux through the plant, determined from the change in pot weight between dawn and evening, prior to the onset of nighttime fertigation. **Whole-Plant Biomass Gain** : the daily fresh-weight gain of the intact plant, determined by comparing pot weights at full water-holding capacity on two consecutive mornings (04:30), after complete drainage and before the start of transpiration. Since soil, container, and other non-plant mass components remained constant, changes in pot weight reflected net biomass accumulation. Calculations followed Halperin et al. (2017), Dalal et al. (2020), and Shenhar et al. (2025).

These platformed plants provided continuous, reliable traces of plant behavior; we co-installed sensor nodes (hereafter, ‘sensor platforms’, SPs) on the same individuals to evaluate how in-canopy microclimate signals relate to, and can proxy, the functional phenotyping measurements.

### Sensor Platforms (SPs) Validation

Environmental conditions in the WSN were monitored using the CC2650F128 SensorTag Kit platform manufactured by Texas Instruments, United States, a compact wireless sensor node integrating multiple environmental sensors, the Sensors Platform (SP). According to manufacturer specifications, the relative humidity (RH) sensor has an accuracy of ±3% and the temperature sensor ±0.2 °C, across a range of 0–100% RH and –40 to 125 °C. The ambient light sensor provides a linear response over a dynamic range of 0.01 to 83,000 lux, with spectral responsivity across the 400–700 nm (Texas Instruments, 2016). Click or tap here to enter text.To complement the manufacturer-provided specifications and ensure sensor performance under our experimental conditions, we conducted independent validation experiments, which confirmed the high accuracy and repeatability of the sensors (R² = 0.997–1.000), as detailed below. Three separate validation experiments were conducted. For temperature and humidity, SP sensors were compared to the LI-COR 6800 Infrared Gas Analyzer (IRGA) and thermometer, using the small plant chamber (6800-17), which allowed the insertion of three SPs per trial (Figure Suppl. 1A). The internal volume of the chamber was 193.2 cm³, with built-in specifications of ±0.15 °C for temperature and ±1.5% for relative humidity (Figure Suppl. 1B–C). For light-intensity measurements, a separate experiment was conducted using the LI-COR 6400-02B LED light-source chamber, configured to emit 15% blue and 85% red light. Three SPs were positioned alongside the reference device, simulating the position of a leaf inside the chamber, and readings were taken at each light level. Measurements were conducted across a light-intensity range of 50–2000 µmol m⁻² s⁻¹ PPFD (corresponding to 3,000–82,000 lux), measured once at each light level (Figure Suppl. 1D).

### Experimental Design

The experimental design was developed to test the hypothesis that real-time monitoring of plant– microclimate interactions can serve as a proxy for physiological responses. Thus, in each experimental run, all plants were grown under similar controlled conditions, including uniform irrigation and climate settings. Planting density and cultivation settings mirrored commercial practice, providing a realistic context for diagnosing plant status from canopy–ambient microclimate differentials. A subset of representative plants was equipped with multi-level microclimate sensors positioned within the canopy (Figure 1). This design enabled comparison between in-canopy and ambient environments, allowing us to evaluate the effects of plant presence and activity on local microclimate and to correlate these environmental changes with plant physiological performance. Each selected plant was equipped with at least one sensor node positioned at the lower canopy (∼30 cm above the pot base). A subset of these plants also had additional sensors at mid-canopy (∼60 cm) and upper canopy (∼90 cm) heights, enabling vertical profiling of temperature, relative humidity, and light intensity within the canopy (Figure 1A). In total, 15 wireless sensor nodes were installed on plants: seven at 30 cm, four at 60 cm, and four at 90 cm; two additional nodes (SP16, SP17) were placed in open areas to serve as ambient references (Figure 1H). This configuration enabled real-time monitoring of plant-driven microclimate modification and assessment of interaction consistency across plants with differing canopy architectures. To evaluate microclimatic differences between the in-canopy and ambient reference environments, we deployed paired sensors per variable, with one inside the canopy (IC) and one at a nearby ambient reference (R). All sensors logged continuously, providing uninterrupted microclimate time series. For analysis, we summarized plant behavior using a representative midday window, selected after the room stabilized following lights-on and the temperature ramp (see Methods and Fig.2A). At each timestamp, differentials were computed as in-canopy minus ambient reference, with 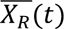 defined as the timestamp-aligned mean of the Watchdog weather station and the two ambient SPs (SP16, SP17) located outside the canopy. To remove pair-specific fixed measurement offsets between the IC and R sensors (which can arise from small calibration tolerances), we computed a pre-dawn technical differential (05:00–06:00) for each IC–reference pair and subtracted it from the midday differential (11:00–12:00), yielding the environmental signal (Equation 1). We selected 11:00–12:00 to coincide with “peak” plant activity, higher radiative loading, and larger gradients, and 05:00–06:00 as a “quiet” baseline period in the room (lights off, minimal lamp heating, negligible plant-driven fluxes), during which cross-node variability is lowest, as indicated by the coefficient of variation (CV) across all SPs (Supplementary Fig. 2).

For all ΔX computations, the ambient reference 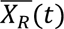 was defined as the timestamp-aligned mean of the WatchDog weather station and the two ambient sensor platforms (SP16 and SP17) located outside the canopy. Equation 1 was applied uniformly to temperature, relative humidity, and light (illuminance), where *X* denotes the variable under comparison in time *t*.

All sensor measurements, including whole-plant transpiration, plant weight, in-canopy microclimate, and weather-station data, were time-synchronized and recorded continuously throughout the growth period of each experimental run. A logging interval of 3 minutes was used for all systems to ensure adequate temporal resolution for capturing rapid plant– environment interactions while limiting data redundancy and noise. This interval was based on prior work demonstrating that 3-minute sampling reliably captures short-term physiological oscillations without information loss (Wallach et al., 2010). The resulting dataset provided a four-dimensional view of plant–environment interactions, incorporating spatial position (x, y, z within the canopy) and time. This allowed dynamic tracking of microclimate gradients and their progression as the canopy developed. All data streams were transmitted in real time to a cloud-connected server, enabling remote monitoring and preliminary data analysis during the experiment.

### Wireless Sensor Network Design and Energy Optimization

To support long-term, energy-efficient telemetry, the CC2650F128SP child nodes were configured to operate in ultra–low-power mode, utilizing only their onboard sensors for ambient temperature (°C), relative humidity (%RH), illuminance (lux), and barometric pressure (mPa) (Texas Instruments, 2016). A CC2650 LaunchPad™ (LAUNCHXL-CC2650 Development Kit TI.Com) acted as the local data-forwarder and radio bridge to a 6TiSCH Border Router (6BR), which maintained network time and relayed traffic to the server on a Raspberry Pi 3B+.

Communication used IEEE 802.15.4e Time-Slotted Channel Hopping (TSCH) with autonomous scheduling function (Orchestra), and the IETF 6TiSCH stack for low-power IPv6 routing (TSCH/6LoWPAN/RPL/UDP). This combination provided reliable delivery at very low energy cost in dense plantings (Texas Instruments, 2016).

For readers less familiar with networking: a small base unit in the room keeps the network’s clock and bridges data to the server (the 6TiSCH Border Router, 6BR). Sensor nodes wake for brief, pre-assigned moments to send/receive and otherwise sleep, while the radio channel changes each moment to avoid interference (time-slotted operation with channel hopping; IEEE 802.15.4e TSCH with autonomous scheduling via Orchestra). Internet headers are compressed so tiny packets fit the low-power link (6LoWPAN); if a node is far, the data can hop via neighbors (multi-hop routing with RPL); and a minimal transport protocol is used to keep overhead low (UDP). The practical upshot is stable delivery with minimal battery drain, even when many plants are instrumented. In practice, a single base unit comfortably supports on the order of ∼100 SPs at our reporting rate; with sparser reporting, capacity increases (consistent with large 6TiSCH testbeds at a few hundred nodes; Duquennoy et al., 2017; Vilajosana et al., 2020; Thubert, 2021).

With the architecture and capacity established, we next quantified how the medium-access policy governs reliability and energy, contrasting contention-based CSMA with the scheduled, channel-hopping TSCH used in our deployments. Under otherwise identical sampling rates and payloads, each SP was instrumented with the Contiki-NG Energest module; we analyzed fixed 60-s windows (“periods”) matching the reporting interval with per-state accounting for CPU, Radio Rx/Tx, Low-Power Mode (LPM), and Deep-LPM (Dunkels et al., 2007; Sabovic et al., 2020). Relative to CSMA, where nodes contend for an apparently free channel, incurring collisions, retries, and idle listening, TSCH assigns transmissions to fixed slots and hops among radio channels, so radios wake only for scheduled slots and otherwise sleep, yielding duty cycles <1% (radios off >99%) and higher stability (Dunkels et al., 2004; 2011; Duquennoy et al., 2017; Mavromatis et al., 2016; Alves and Zhai, 2017). Consequently, energy spent on radio reception dropped sharply under TSCH despite a modest rise in CPU time, producing lower total consumption (Supplementary Figure 3). Battery-lifetime estimates derived from these profiles indicated ≈120 days per SP under TSCH versus ≈11 days under CSMA (Supplementary Figure 4). In topology tests (1/2/8 nodes spanning 0.87–6.86 m from the border router), cumulative-energy slopes stabilized after ∼60 periods (6 h runs of 360 periods), and lifespans declined only modestly with distance, consistent with slightly higher communication load (Supplementary Figure 5). In a growth-room deployment of 17 SPs (arranged as in Figure 1), observed lifetimes averaged 109 ± 6.3 days per SP, consistent with model predictions.

### Packet Loss Measurements

To evaluate transmission reliability under conditions more demanding than the growth room (closer to operational agriculture in terms of microclimate variability and radio propagation) we conducted two deployments in semi-controlled greenhouses at the I-CORE Center: a polycarbonate dual-gable structure (18 × 16 m) and a single-span glasshouse (10 × 5 m), both located on the same campus. Data integrity was assessed by analyzing packet loss and gap characteristics, where a gap was defined as a continuous sequence of missing 3-minute intervals (‘NaN’ values) in the dataset timeline. Each SP was programmed to transmit duplicate data packets twice every 3 min to introduce redundancy and improve delivery reliability; only the latest received packet within each interval was stored in the cloud database. The polycarbonate greenhouse deployment included 144 SPs operating for 11 days, resulting in 755,000 expected data points. The glasshouse deployment included 64 SPs operating for 30 days, generating 983,000 expected data points. To compute packet loss, an independent timestamp array was generated for each SP, spanning from its first to last recorded timestamp, assuming one packet every 3 min. Missing timestamps were replaced with ‘NaN’ values, producing a complete reference timeline. To ensure validity, two filtering criteria were applied: (1) gaps longer than two hours were excluded, as these represent hardware-level failures (e.g., power or battery issues); and (2) sequences of three or more identical consecutive values were considered sensor failures, with all repeated values beyond the third replaced by ‘NaNs’ until a valid change occurred. Packet loss for each SP was calculated as the percentage of missing timestamps relative to the total expected 3-minute intervals. Gap duration was defined as the number of consecutive ‘NaN’ intervals, each representing 3 minutes of missing data. These metrics were used to characterize both packet transmission quality and the duration of communication interruptions (Supplementary Figure 6). To isolate the effect of redundancy, we repeated the analysis in single-transmission mode, one packet every 3 min with no duplicate retransmission, using the same reporting interval, topology, and filtering rules as above. Two deployments were analyzed: 66 sensor nodes for 60 days in the growth chamber, and 64 sensor nodes for 90 days under the same configuration. Under this single-transmission mode, packet loss (%) and gap durations increased, demonstrating the contribution of redundancy to communication reliability (Supplementary Fig. 7). **In summary**, protocol evaluation identified TSCH as the most suitable solution for high-frequency plant–environment monitoring, providing an optimal balance between temporal resolution, energy efficiency, and transmission reliability (Dunkels et al., 2004; 2011; Duquennoy et al., 2016; Mavromatis et al., 2016; Alves and Margi, 2017). Redundant dual transmissions further reduced packet loss and communication gaps under greenhouse conditions; while this approach carries an energy cost, we prioritized data completeness and therefore continued operating with dual transmissions.

### Pricing

One of the key advantages of the WSN system is its ability to provide dense spatiotemporal coverage by deploying multiple sensor nodes operating continuously and in parallel. While such resolution might suggest high cost due to the number of spatially distributed, constantly transmitting units, the system remains relatively inexpensive. For example, the deployment used in this study included 17 Sensor Platforms (SPs; $60 each) together with a Raspberry Pi 3B ($80) and a CC2650 LaunchPad ($15), resulting in a total hardware cost of approximately $1,115. This configuration demonstrates a scalable and cost-effective solution for real-time, high-resolution environmental monitoring in agricultural systems. In practical terms, this setup functions as a 17-node array of continuously streaming micro-meteorological stations (T, RH, illuminance, pressure), delivering dense, user-defined spatial coverage at field-relevant timescales. Additional coverage scales linearly at ≈$60 per node (excluding the one-time base unit), yielding a per-instrumented-plant cost of ∼$66 in the present deployment.

Together, these methods enabled simultaneous, high-resolution environmental and physiological monitoring, linking WSN-derived microclimate data with whole-plant water use and growth dynamics, and providing an integrated framework for evaluating plant–environment interactions.

## Results

After verifying sensor accuracy (Supplementary Figure 1), we confirmed that the WSN met all predefined requirements for high spatiotemporal plant–environment monitoring. TSCH communication significantly reduced energy consumption and extended battery life compared to CSMA (Supplementary Figures 3-4). Across topologies, estimated battery life decreased by <15% with added hops or distance to the border router (Supplementary Fig. 5; Methods), consistent with modest increases in radio duty cycle; critically, runs still spanned few months. Accordingly, topology can be chosen primarily for sensing objectives and placement logistics rather than energy constraints; even where battery replacement becomes slightly more frequent, a compromise we preferred to make to maintain continuous, high-frequency telemetry.

Dual-transmission mode ensured low packet loss (1.5–3.7%) and short data gaps, whereas single transmission increased both metrics (Supplementary Figures 6–7). This redundancy carries only a modest energy overhead: under TSCH the dominant radio cost is reception rather than transmission, so duplicating short packets adds little to total consumption (see the TSCH energy breakdown in Supplementary Figure 3). Consistent with Energest-based projections and topology tests (Supplementary Figures 4–5), network lifetimes remained ∼3 months across practical layouts. We therefore adopted dual transmission as the default, prioritizing data integrity, while noting that redundancy can be throttled adaptively where energy budgets are paramount (e.g., seasonally or by node–router distance).

With this validated infrastructure, we next tested our central hypothesis that in-canopy versus ambient microclimate differences can serve as real-time indicators of plant physiological status. First, we continuously monitored environmental conditions and plant growth within the controlled growth room. Measurements from the WatchDog weather station showed stable photoperiod cycles (Figure 2A) and consistent diurnal fluctuations in relative humidity (Figure 2B) and air temperature (Figure 2C). As a representative example, plant biomass, derived from the PlantArray system, increased steadily throughout the experiment (Figure 2D), reflecting continuous vegetative development under stable environmental conditions.

The subsequent phase of our analysis focused on validating the reliability of the sensor platforms (SPs) for environmental monitoring. For this purpose, two reference SPs were deployed at distant locations within the growth room: SP16 adjacent to the WatchDog weather station and SP17 positioned on the room periphery, both outside the immediate influence of plant canopies (see Figure 1 and Methods). These SPs provided a basis for comparing WSN-derived measurements with those from the standard weather station. Temperature data from the two reference SPs showed good correlation with the WatchDog readings (Figure 2E) and exhibited similar diurnal patterns over a representative five-days period (Figure 2F). While minor offsets were observed, likely resulting from local microclimatic differences between sensor locations, the overall agreement confirmed the accuracy and stability of the SP measurements. This validation provides the basis for using the WSN to track plant-driven modifications of their immediate environment, forming the foundation for the subsequent analyses addressing our central research hypothesis. Accordingly, for all ΔX computations (Eq. 1) we defined the ambient reference 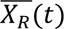 as the timestamp-aligned mean of the WatchDog station and the two ambient nodes (SP16, SP17), since sensitivity checks showed that normalizing with any single device or pairwise combination yielded closely similar differentials and downstream relationships.

To evaluate whether the SP-based WSN can capture physiological responses relevant to plant growth, we focused on seven plants that were simultaneously monitored by SPs for microclimate parameters and by the PlantArray system for physiological traits (Figure 1). Biomass accumulation in these plants increased steadily during the one-month monitoring period (Figure 3A), accompanied by an increase in daily transpiration (Figure 3B). When averaged across all plants, biomass gain and daily transpiration followed highly similar temporal patterns (Figure 3C). A correlation analysis revealed a strong positive relationship between daily transpiration and biomass gain (R² > 0.9; Figure 3D), confirming that whole-plant water use is closely linked to growth under these controlled conditions.

**Figure 3.**
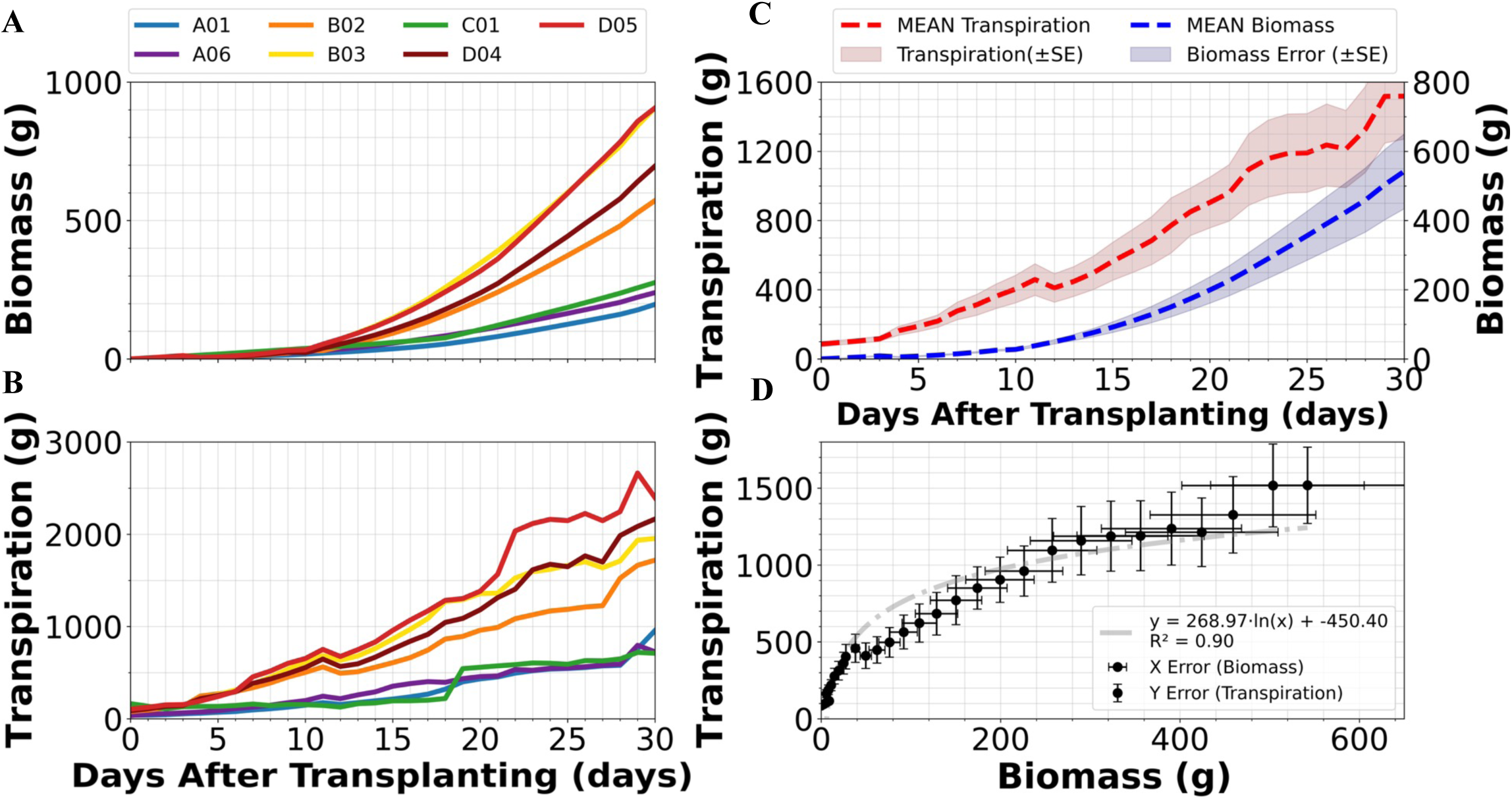
Biomass gain and whole-canopy transpiration of PlantArray-monitored plants that were simultaneously monitored by SP sensor platforms. (A) Net biomass gain (g) of seven *Cannabis sativa* (MVA strain) plants (A01, A06, B02, B03, C01, D04, D05; see Fig. 1 and Methods), monitored using the PlantArray functional phenotyping system. These same plants were also continuously monitored by SPs for microclimate measurements. (B) Daily transpiration (g day⁻¹) of the same plants, showing increasing water use over time. (C) Combined time-series of mean biomass gain and mean transpiration, demonstrating parallel growth dynamics where transpiration changes closely followed biomass accumulation. (D) Correlation between daily transpiration and biomass gain across all seven plants over the entire monitoring period (mean ± SE). The strong positive relationship (R² > 0.9) highlights whole-plant transpiration as a robust indicator of biomass gain under these controlled conditions.

Using the canopy–reference pairs (described in Methods), midday measurements revealed that the canopy cooled its immediate environment relative to ambient reference conditions, with mean differences of ∼4 °C near the canopy base (30 cm) and ∼7 °C near the upper canopy (60 cm; Figure 4A, B). These differences decreased gradually during development, reaching a stable cooling pattern once canopy closure over SP (at given height) was achieved. When canopy temperature values were plotted against plant biomass, the relationship exhibited a negative logarithmic saturation trend (Figure 4C), demonstrating that canopy cooling capacity rapidly increased during early growth and then stabilized as plant biomass increased.

**Figure 4.**
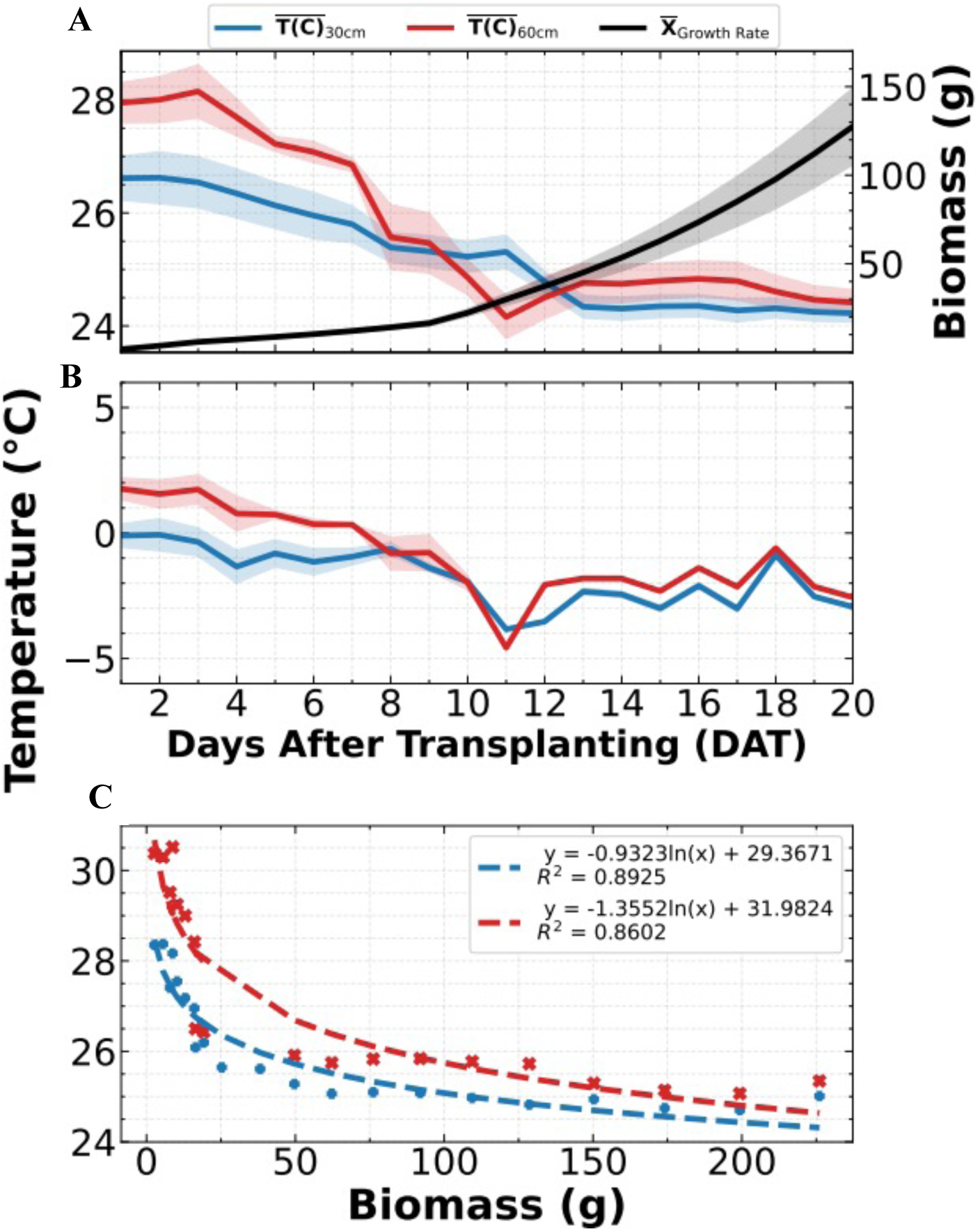
In-canopy temperature dynamics at two canopy heights relative to a reference sensor and their relationship to plant growth. (A) Mean midday in-canopy temperature (°C; left axis) measured at 30 cm (blue) and 60 cm (red) canopy heights, plotted daily for the first 20 days after transplanting (DAT). Shaded areas represent ±SE across seven *Cannabis sativa* seedlings (A01, A06, B02, B03, C01, D04, D05; see Fig. 3 and Methods). Calculated plant biomass (g; right axis) is shown for the same period.(B) Mean temperature difference (ΔT) between each canopy sensor and its paired reference sensor, calculated using Equation 1 (see Methods). Shaded areas represent ±SE, which when not visible are smaller than the line thickness. The data show that canopy microclimate cooling increased as plants developed, with a stable equilibrium reached at ∼4 °C cooling near the canopy base (30 cm) and ∼7 °C near the upper canopy (60 cm).(C) Relationship between canopy temperature and plant biomass at each height. Logarithmic regression fits indicate a saturation trend, where microclimate cooling effects are strongest at early growth stages and stabilize as the canopy reaches maturity.

Note: Data from the 90 cm SPs were excluded from the analysis because the plants never reached this height during the experimental period, and the lamp-raising method caused anomalous and inconsistent readings.

Similar to the canopy cooling effect observed in temperature measurements, we also detected a strong relationship between plant biomass and the reduction of in-canopy light intensity. Illuminance inside the canopy declined consistently as plants developed (Figure 5A,B). Notably, the lower canopy (30 cm) responded first, showing an earlier reduction in light penetration and reaching a steady attenuation level more quickly. In contrast, the upper canopy (60 cm) exhibited a delayed but pronounced reduction, consistent with the timing of stem elongation and canopy expansion at higher levels (Figure 1C,D). As plant biomass increased, these vertical differences diminished, eventually reaching a stable attenuation pattern once canopy closure was achieved. When in-canopy illuminance was plotted against biomass, both canopy heights followed an inverse logarithmic saturation trend (Figure 5C). This indicates that self-shading intensifies rapidly during early growth stages and then stabilizes as canopy structure matures. Notably, these measurements, recorded in lux, are strongly correlated with photosynthetically active radiation (PPFD) equivalents, based on the linear relationship between SP sensor readings and a LI-COR LI-250A spectrometer (Supplementary Figure 8).

**Figure 5.**
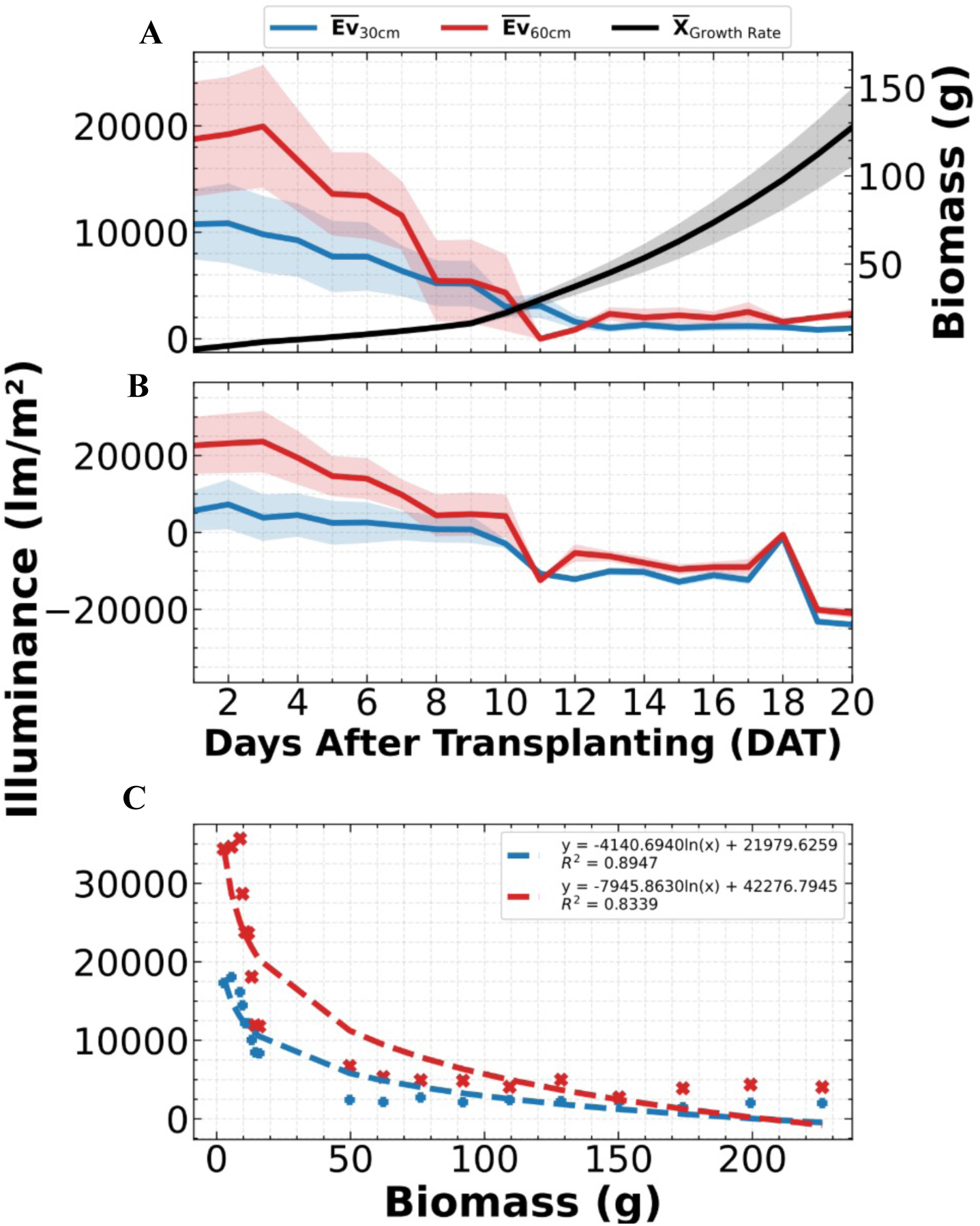
In-canopy light intensity dynamics at two canopy heights relative to a reference sensor and their relationship to plant growth. (A) Mean midday in-canopy illuminance (Lux, lm m⁻²; left axis) measured at 30 cm (blue) and 60 cm (red) above the pot base, plotted daily for the first 20 days after transplanting (DAT). Plant biomass (g; right axis) is shown for the same period (same plants as in Figure 4). (B) Mean illuminance difference between each canopy sensor and its paired reference sensor, calculated using Equation 1 (see Methods). Shaded areas represent ±SE; where not visible, SE is smaller than the line thickness. The lower canopy (30 cm) exhibited an earlier reduction in light penetration, reaching equilibrium sooner, whereas the upper canopy (60 cm) responded later, consistent with stem elongation and canopy expansion. (C) Relationship between in-canopy illuminance and plant biomass for both canopy heights. Logarithmic regression fits indicate a saturation trend, where light attenuation increases rapidly during early growth stages and then stabilizes as canopy structure matures.

After examining the effects of biomass growth on canopy temperature and light attenuation, we next investigated how plant transpiration influences in-canopy relative humidity (RH). Midday RH, measured at 30 cm and 60 cm above the pot base, increased progressively as daily whole-plant transpiration rose (Figure 6A), stabilizing as leaf area expanded. This absolute increase in in-canopy RH (18–25% higher than the paired reference sensors, depending on canopy height; Figure 6B) indicates intensive plant activity driven primarily by increasing transpiration rates and reflects both the vertical progression of canopy development and the gradual establishment of humidity gradients within the plant microclimate. When RH was plotted against daily transpiration, both canopy heights followed a logarithmic saturation trend (Figure 6C). This indicates that increases in transpiration rapidly elevate in-canopy humidity during early growth but reach an equilibrium once canopy structure and leaf conductance stabilize.

**Figure 6.**
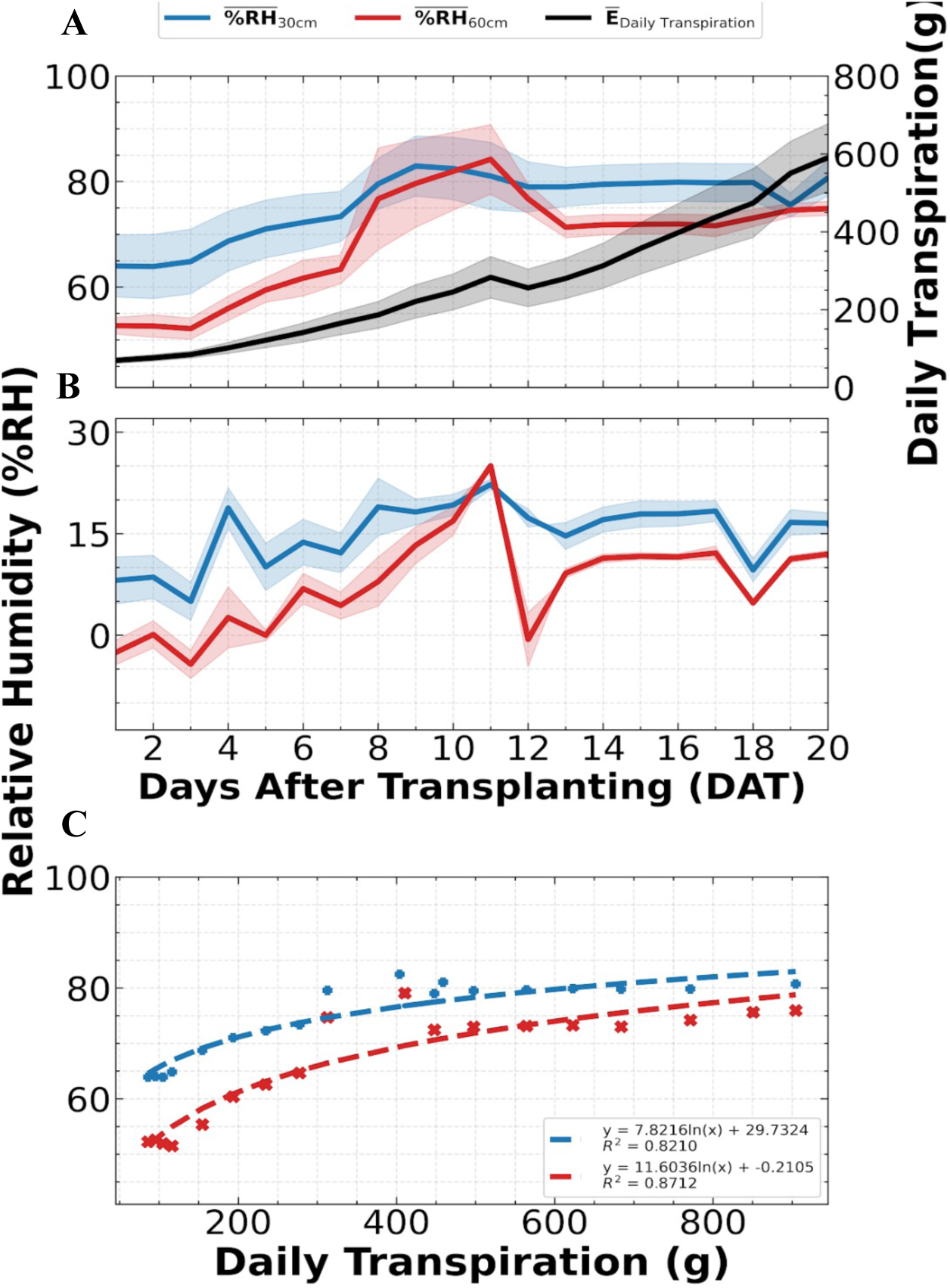
In-canopy relative humidity (RH) dynamics and their relationship to plant transpiration. (A) Mean midday in-canopy RH (%) at 30 cm (blue) and 60 cm (red) above the pot base, plotted for the first 20 days after transplanting (DAT), with daily whole-plant transpiration (g; black) shown on the right axis (same plants as in Figure 4). (B) Mean RH difference between each canopy sensor and its paired reference sensor, calculated using Equation 1 (see Methods). Shaded areas represent ± SE; where not visible, SE is smaller than the line thickness. Both canopy heights showed a positive RH differential that increased as transpiration rose, with a stronger early response at 30 cm and delayed response at 60 cm. (C) Relationship between daily transpiration and in-canopy RH at both heights, showing logarithmic saturation trends (dashed lines) indicating that RH increases with transpiration but approaches a stable canopy microclimate at high transpiration rates.

Repeating these experiments with the bigger canopy “Sativa-like” Odem cultivar (Figure 1F) using an identical experimental design produced plant–environment interaction patterns closely matching those observed for the MVA cultivar (Figure 7). The canopy again imposed a vertically structured microclimate. In-canopy air temperature decreased as biomass accumulated (Figure 7A–B), consistent with progressive canopy cooling as foliage expanded (logarithmic fits, R² ≈ 0.91–0.95). Illuminance within the canopy declined with biomass (Figure 7C–D), following logarithmic attenuation (R² ≈ 0.82–0.91). Relative humidity increased in parallel with daily transpiration (Figure 7E–F), with saturating logarithmic relationships (R² ≈ 0.97–0.98).

**Figure 7.**
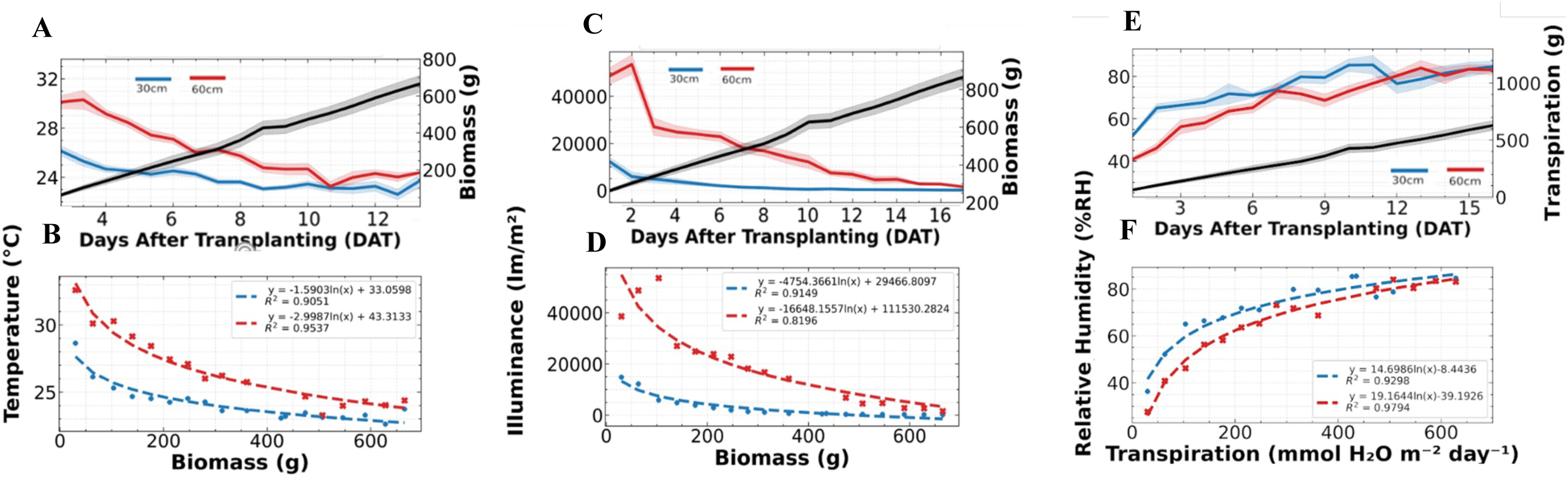
Plant microclimate dynamics in Cannabis sativa ‘Odem’ (Sativa-like morphology). (A) Mean midday in-canopy air temperature (°C) at 30 cm (blue) and 60 cm (red) with biomass (black, right axis) over days after transplanting (DAT). (B) Relationship between in-canopy temperature and plant biomass (g), with logarithmic cooling fits. (C) Mean midday in-canopy illuminance (lux; 1 lux = 1 lm m⁻²) at 30 cm (blue) and 60 cm (red) with biomass (black, right axis). (D) Relationship between in-canopy illuminance and biomass, with logarithmic attenuation fits. (E) Mean midday in-canopy relative humidity (%RH) at 30 cm (blue) and 60 cm (red) with daily transpiration (black, right axis). (F) Relationship between in-canopy RH and plant transpiration (mmol H₂O m⁻² day⁻¹), with logarithmic fits. Shaded areas denote mean ± SE; where not visible, SE is smaller than the line thickness.

Overall, the early and stronger microclimatic shifts at the lower canopy, followed by slower but steady responses at the upper canopy, were consistent between the compact MVA and the more open, Sativa-like Odem cultivar.

## Discussion

Our results demonstrate that plant-driven microclimatic shifts (cooling, humidification, and light attenuation within the canopy) can be captured continuously by a low-cost WSN, and that these canopy–ambient differentials scale with transpiration and biomass across contrasting architectures. Accordingly, we next consider how our approach compares with existing sensing technologies and what its implications are for large-scale, continuous phenotyping.

As a whole, our findings show that a standards-based, canopy-integrated WSN can resolve plant-driven microclimatic signals continuously and with engineering robustness, positioning canopy– ambient differentials as operational proxies of physiological status. The telemetry layer exhibited low packet loss, months-long operation, and strong agreement with independent references (Figures 2E–F; Supplementary Figures 1, 3–7), indicating that the observed dynamics reflect biology rather than measurement artifacts. Under stable conditions, daily transpiration tightly predicted biomass gain (R² > 0.9; Figure 3D; Shenhar et al., 2025; Dalal et al., 2019), and the same process was mirrored in the microclimate: progressive evaporative cooling (lower in-canopy temperature), humidification (higher in-canopy RH), and optical attenuation (declining in-canopy illuminance) followed saturating logarithmic relationships with growth or transpiration, with earlier and stronger responses in the lower canopy (Figures 4–6). Repeating the experiment in the structurally distinct ‘Odem’ cultivar reproduced these patterns (Figure 7), supporting robustness across contrasting architectures and reinforcing the generality of the WSN approach.

Collectively, these validations justify treating the WSN as an always-on, canopy-integrated measurement layer for canopy–ambient differentials, suitable for real-time inference of plant status in operational settings. In practice, microclimate differentials function as deployable physiological proxies that complement direct instruments and imaging, offering a scalable, lower-cost route to continuous phenotyping and early stress diagnostics at field scale.

Using physiological parameters as proxies for plant behavior, development, and stress responses is well established (Zimmermann et al., 2013; Negrão et al., 2017; Dalal et al., 2019). Its chief advantage is diagnostic lead time: changes in stomatal conductance and whole-plant transpiration emerge within minutes to hours after a perturbation, well before visible symptoms such as wilting or chlorosis, when damage is still reversible and interventions (e.g., irrigation or fertigation) are most effective. However, the high-precision instruments typically used for such measurements are costly, technically demanding, and low-throughput (e.g. leaf-level gas-exchange systems, pressure-chamber measurements of leaf water potential, and sap-flow/psychrometric probes), limiting season-long or field-scale deployment and increasing reliance on skilled scouting, which limits season-long or field-scale deployment and increases reliance on skilled scouting. These constraints motivate scalable surrogates that retain physiology’s early-warning benefit while minimizing cost and operator burden. In addition to direct physiological diagnosis, a critical element of our framework is the fourth dimension— time, which furnishes an internal reference for each plant and enables cohort-level comparison of developmental trajectories against concurrently monitored peers (and, prospectively, against site-specific environmental references). In this setting, continuously streaming, canopy-integrated microclimate differentials (ΔT, ΔRH, ΔLight) provide per-plant baselines and cohort benchmarks: deviations from a plant’s own temporal trajectory or from its peers flag emerging developmental or stress-related divergence. Accordingly, specific, testable signatures in ΔT, ΔRH, and ΔLight can serve as early indicators of stomatal constraints (e.g., water stress) and support timely intervention. Looking ahead, such per-plant temporal baselines could underpin adaptive, cultivar- and site-specific thresholds, benchmarking across seasons, and calibration of digital-twin models for early-warning decision support.

### Anticipated microclimatic signatures under water stress and other stomatal constraints

Diverse abiotic and biotic stressors often trigger rapid stomatal closure, reducing transpiration within minutes to hours. Accordingly, we expect rapid microclimatic signatures: (i) canopy cooling weakens as the temperature differential 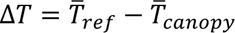 contracts toward zero and may invert at midday under matched external irradiance; (ii) the humidity differential 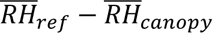 declines relative to plant-specific baselines; and (iii) in-canopy light attenuation relaxes (ΔLight decreases) as leaf angle/rolling increases light penetration. Over hours to days, these fast responses propagate to daily integrals, reduced daily transpiration, slower biomass accumulation, diminished self-shading, and damped diurnal amplitudes of all differentials. Upon stress end (e.g. re-watering), recovery kinetics (e.g., nighttime rehydration and the rebound rate of transpiration) yield quantitative resilience metrics. These expectations align with established physiology in salinity/drought and with high-throughput functional phenotyping (Zimmermann et al., 2013; Negrão et al., 2017; Dalal et al., 2019).

Continuous dual-stream monitoring of in-canopy and ambient microclimate thus exposes both immediate physiological responses and slower developmental trends under operational conditions, particularly valuable for field phenotyping where in situ, high-frequency dynamics are otherwise hard to capture. The resulting 4D traces pair naturally with emerging digital-twin and machine-learning workflows that depend on dense, labeled streams for synchronization, calibration, and forecasting (Escribà-Gelonch et al., 2024; Zhang et al., 2025; Zamora-Izquierdo et al., 2019; Benos et al., 2021; Zhai et al., 2020).

Taken together, because many abiotic and biotic stressors converge on stomatal regulation, often provoking rapid closure and reduced transpiration (Fridman et al., 2025), we do not expect our microclimate-based assessment to discriminate etiologies in its initial form. Instead, its primary value is cause-agnostic early warning: sustained departures of ΔT, ΔRH, and ΔLight from plant-specific temporal baselines and cohort norms flag deteriorating status and prompt targeted follow-up. By functioning as an always-on, canopy-integrated sentinel, the WSN shortens time-to-detection and expands the intervention window, the practical gain we prioritize at this stage.

### Economic considerations and scale

Beyond technical performance, the economic profile of a canopy-integrated WSN favors scale. Each sensing point is reusable across seasons, so marginal costs decrease as networks grow. Operating costs are dominated by scheduled battery service, while data backhaul and storage are lightweight due to small, infrequent packets. In contrast, high-precision physiological instruments (e.g., IRGA systems, pressure chambers, sap-flow/psychrometric probes) are capital-intensive, require skilled operators, and remain low-throughput. Imaging-based phenotyping (UAV/airborne or fixed arrays) entails specialized operation, regulatory/mission overhead, and substantial processing pipelines, and typically offers discrete temporal “snapshots.” Consequently, on a per-plant basis the WSN provides substantially lower cost for continuous, labeled time series, enabling dense coverage where reference instruments are reserved for calibration and periodic spot-checks. This cost structure supports deployments spanning dozens to hundreds of plants, while complementing remote and proximal imaging in integrated phenotyping workflows.

### Challenges, constraints, and considerations

Our results establish a viable path for continuous, canopy-integrated monitoring, yet several practical constraints should guide interpretation and deployment: Scope and generalizability. Experiments were conducted with two contrasting Cannabis sativa cultivars in a controlled growth room. Although responses were consistent across architectures, broader validation across crops, seasons, and management regimes is required for field-scale generalization.

### Sensor suite and coverage

SPs measured air temperature, RH, and illuminance (lux). While lux was linearly related to PAR in our calibration, it is not spectrally explicit and can saturate near full sun; adding PAR/PPFD, CO₂, leaf-wetness, and wind sensors would expand physiological interpretability. Accuracy, drift, and calibration. Sensor accuracy is high within lab-like ranges, but long-term exposure (dust, condensation, radiation) can induce drift. Our nightly offset correction (IC vs. reference) mitigates systematic bias; periodic cross-calibration and occasional spot-checks with reference instruments are recommended.

### Deployment at scale

Translation from growth room to open fields introduces weatherproofing, installation density, and routine maintenance constraints. Site surveys, height-stratified placement, and adaptive layouts will be necessary to maintain signal quality.

### Connectivity and interference

TSCH with channel hopping and duplicate transmissions reduced packet loss and gaps, but dense networks and complex topologies can exhibit transient route instability that may co-vary with crop phenology. Network health monitoring and conservative duty-cycle budgets are advisable.

### Conclusions

We propose a delta-based, always-on, canopy-integrated phenotyping concept: rather than sensing plants only through episodic imaging or point measurements. We continuously track the canopy–ambient differentials (ΔT, ΔRH, ΔLight) that a plant imposes on its microenvironment, using a lightweight WSN and a simple pre-dawn offset/midday window recipe. In practice, this turns microclimate into a physiological signal layer, streaming at plant resolution, alignable across space and time, and directly comparable within cohorts, thereby elevating environmental telemetry to an operational proxy for plant function. As a system, the approach is modular and scalable. A low-power, standards-based WSN provides months-long, low-loss telemetry at field-relevant densities; the layer can stand alone as a low-cost sentinel or interoperate with existing imaging and functional-phenotyping platforms. Because many stressors converge on stomatal regulation, departures of the differentials from plant-specific temporal baselines yield cause-agnostic early warning, prompting targeted follow-up well before morphological symptoms emerge. Looking ahead, the same streams provide the substrate for adaptive thresholds, edge inference, and digital-twin workflows. Immediate priorities include multi-crop/season pilots, expansion of the sensor suite, modest energy harvesting or scheduled battery service for multi-season runs, and standardized curation pipelines so datasets remain versioned, traceable, and ready for deployment at scale.

## Definitions and Formulas for Environmental Difference Calculation

Let 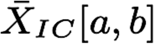 and 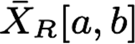 be defined as:

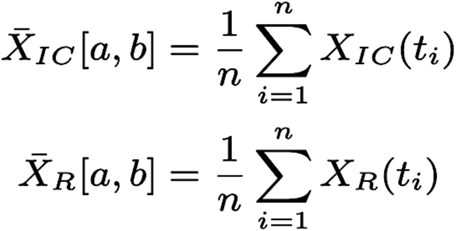

where:

- *X1c(t)* = measurement of variable *X* inside canopy at time *t*
- *XR(t)* = measurement of variable *X* at reference location at time *t*
- *[a, b]* = time interval in hours
- *n* = number of discrete measurements in [*a, b]*

Define:

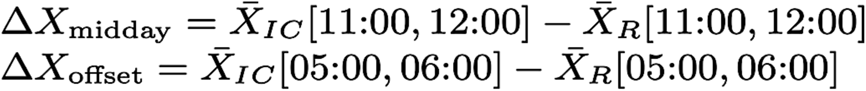

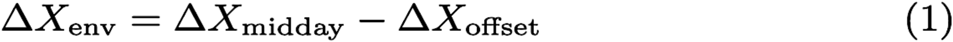

***Equation 1*** *:Notation and formulas for computing environmental difference from canopy and reference measurements, adjusted for technical offset*.

