## Supplementary Figure 1 for "Wireless Sensor Network: New Concept of Spatial-Temporal Monitoring Plant–Environment Interactions"

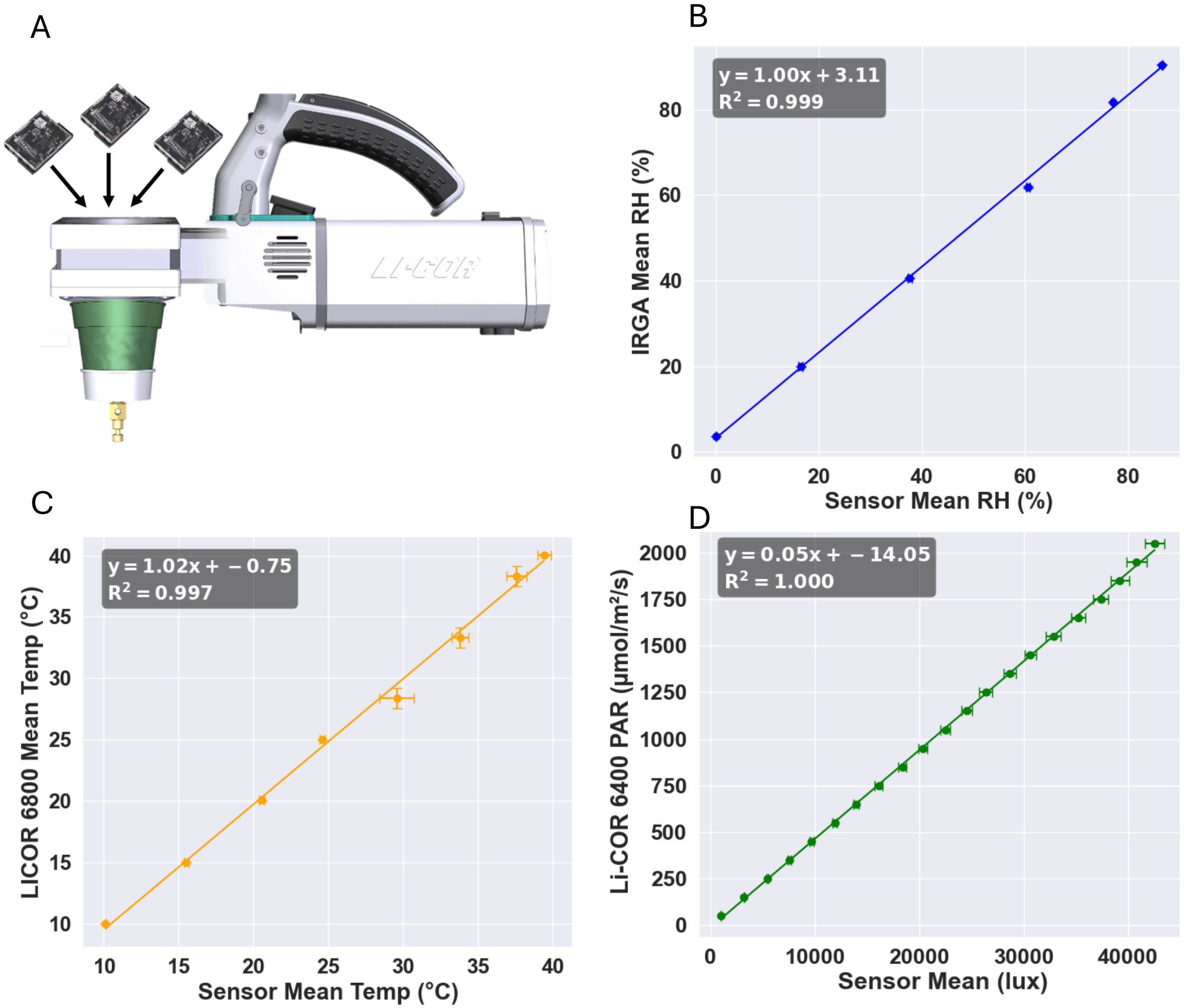


**Supplementary Figure 1. Validation of Sensor Platform (SP) measurements against reference instruments.**
(A) SensorTag units (top left) placed inside the LI-COR 6800 small whole-plant chamber (6800-17) for validation of temperature and relative humidity measurements using the integrated Infra-Red Gas Analyzers (IRGA) and temperature measurement systems.(B) Linear correlation between mean RH values measured by SP sensors and the IRGA reference ($R^2$ = 0.999). (C) Linear correlation between SP-measured temperature and LI-COR 6800 temperature values ($R^2$ = 0.997). (D) Light intensity validation using SP sensors positioned inside the LI-COR 6400-02B LED light source chamber, configured to emit 15% blue and 85% red light. SPs were positioned to simulate the placement of a leaf inside the chamber, and readings were taken simultaneously with the LI-COR 6400 PAR sensor across a range of 50–2000 µmol $m^{-2}$ $s^{-1}$ PAR (corresponding to ~3,000–40,000 lux). The data show a strong linear correlation ($R^2$ = 1.000), confirming high accuracy of the SP light sensors under controlled red-blue spectral conditions.
All data points represent means ± SE of three replicate sensors per light level. Error bars are shown on both axes; where not visible, the standard errors are smaller than the symbol size.
