## Supplementary Figure 2 for "Wireless Sensor Network: New Concept of Spatial-Temporal Monitoring Plant–Environment Interactions"

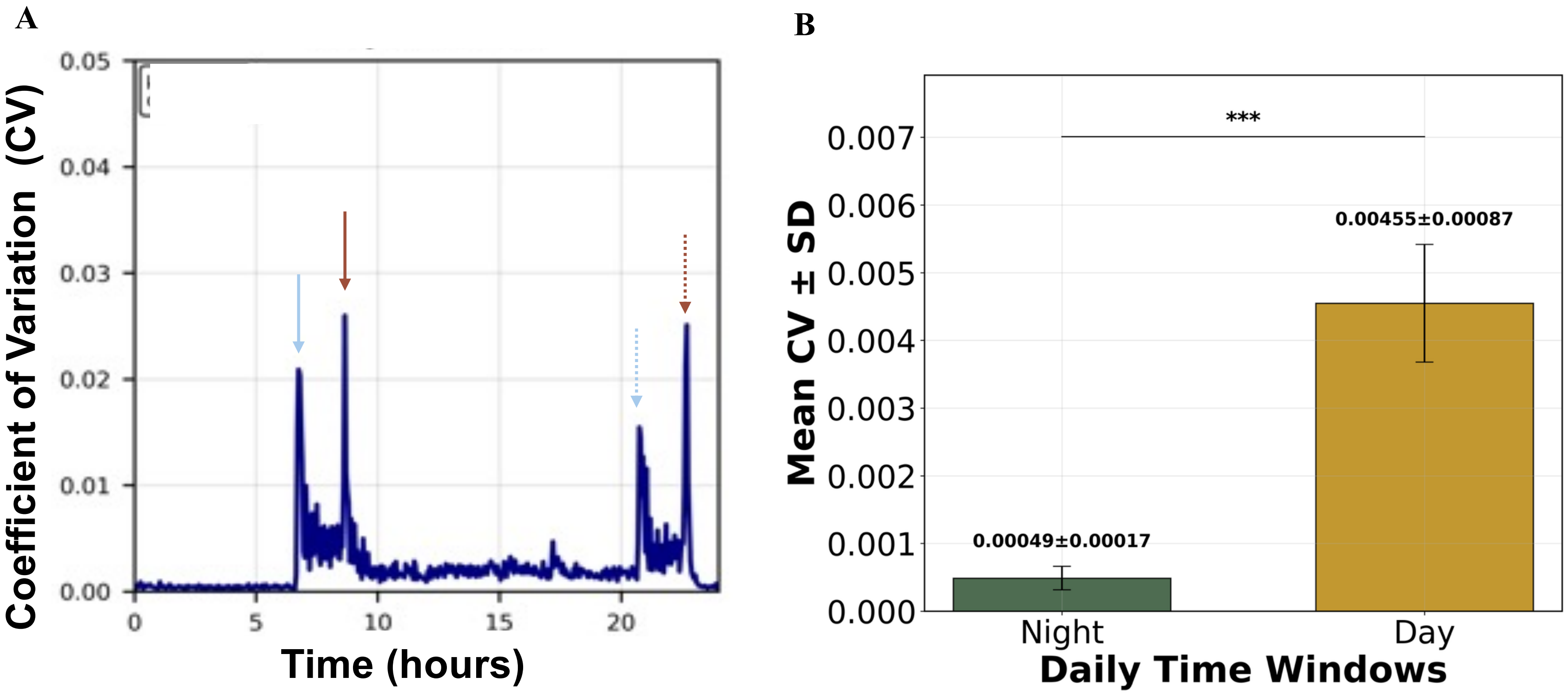


**Supplementary Figure 2. Measurement stability in the growth room under a stepped-spectrum photoperiod.**
(A) Cross-node coefficient of variation (CV = SD/mean; unitless) of air temperature at 3-min resolution over a representative 24-h cycle (16 h light / 8 h dark). Step changes in the lighting program produce brief CV excursions. Arrows mark spectrum events: 4000 K ON (solid light-blue), 4000 K OFF (dotted light-blue), 3000 K ON (solid red), 3000 K OFF (dotted red). CV at night remains low and flat relative to daytime.(B) Mean CV ± SD across 16 days for two analysis windows: Night (05:00–06:00) vs Day (11:00–12:00). Night CV is an order of magnitude lower than Day CV (Welch's t-test, $p < 0.001$), confirming temporal stability during the offset-calibration window. (CV computed per 3-min frame across 17nodes; panel B averages per-day CV across nodes; n = 17nodes × 16 days).
