## Supplementary Figure 3 for "Wireless Sensor Network: New Concept of Spatial-Temporal Monitoring Plant–Environment Interactions"

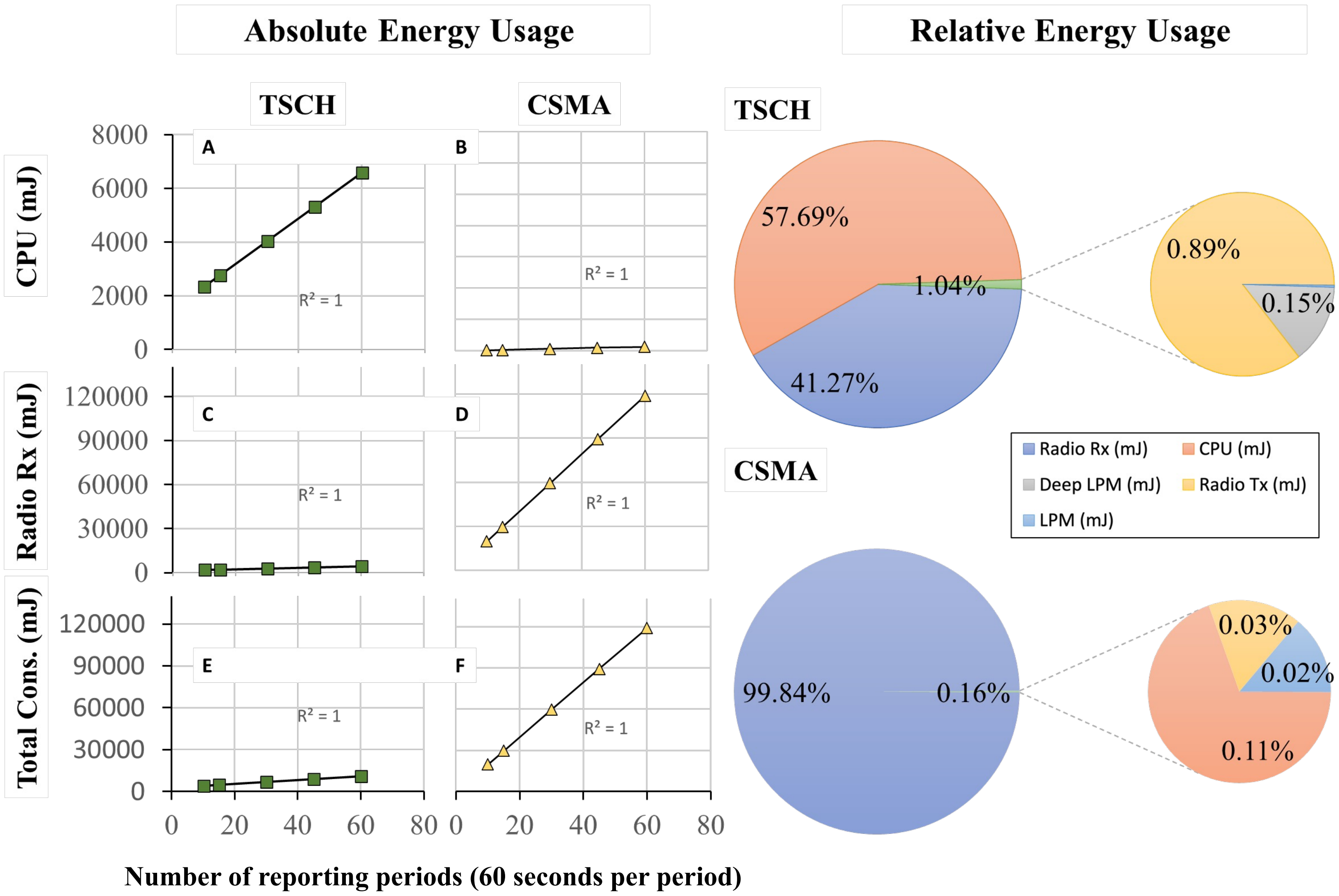


**Supplementary Figure 3.** Comparative energy consumption of two medium-access protocols (TSCH vs. CSMA). Line plots (left) show cumulative energy since $t_0$; the x-axis units are successive 60-s "periods" that match the reporting interval. Absolute energy was measured with the Contiki-NG Energest module for **CPU** (A, B), **Radio Rx** (C, D), and **Total consumption** (E, F). Pie charts (right) summarize the relative share of each operational state for **TSCH (top)** and **CSMA (bottom)** over the same runs: **Radio Rx** — receiver on/listening/decoding (including idle listening in CSMA); **Radio Tx** — transmitter on/sending data and link-layer acknowledgments (MAC ACKs); **CPU**; **LPM** — microcontroller low-power sleep with fast wake; **Deep LPM** — deeper standby/hibernate (most clocks off; slower wake). Under TSCH, 57.7% of total energy was spent in CPU and 41.3% in radio reception, whereas under CSMA, 99.8% was consumed by continuous radio reception, highlighting its much higher baseline power demand.
