## Supplementary Figure 4 for "Wireless Sensor Network: New Concept of Spatial-Temporal Monitoring Plant–Environment Interactions"

**Supplementary Figure 4. Estimation of Battery Lifespan under Different Transmission Protocols**.

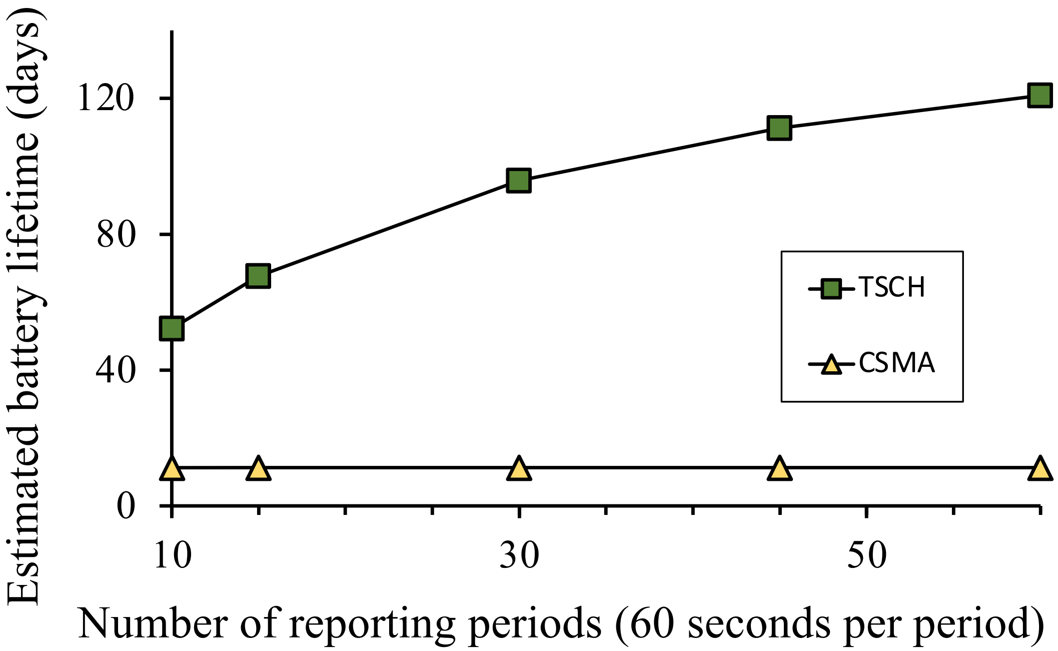


**Supplementary Figure 4. Estimation of Battery Lifespan under Different Transmission Protocols**.
Estimated lifetime of a single Sensor Platform (SP) node as a function of *Energest periods*, where each period represents a 60 s measurement window of energy consumption (single transmission per 60-s period). Results are shown for nodes operating with TSCH (green squares) and CSMA (yellow triangles), illustrating the effect of protocol choice on battery longevity.
