## Supplementary Figure 5 for "Wireless Sensor Network: New Concept of Spatial-Temporal Monitoring Plant–Environment Interactions"

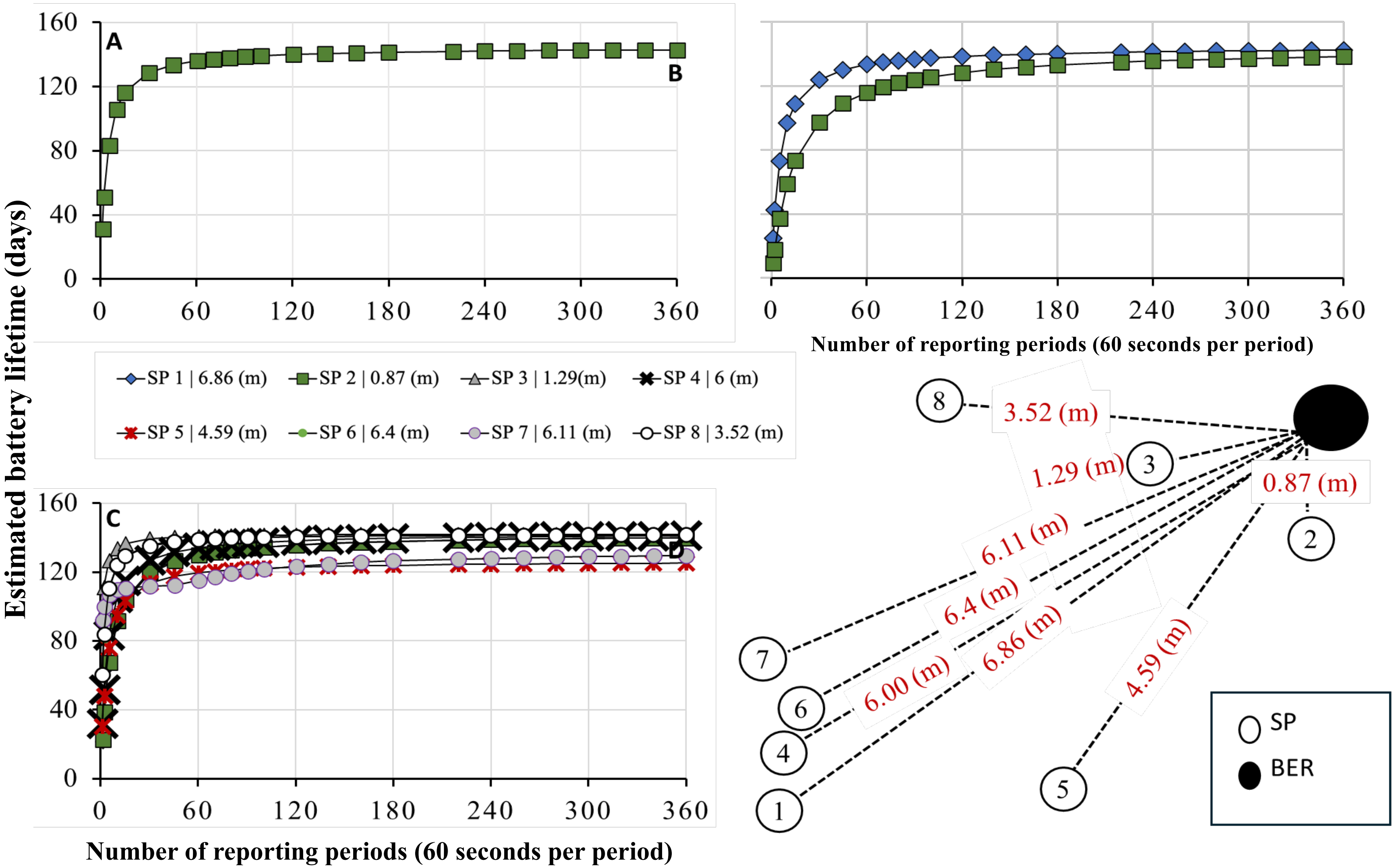


**Supplementary Figure 5. Impact of network topology on battery lifespan.** (A) Lifetime estimation for a single SP node. (B) Two SP nodes are positioned at different distances from the 6TiSCH border router (BER). (C) Eight SP nodes are placed at varied distances. (D) Network layout showing node positions and their relative distances from the BER. Battery lifespan stabilized after approximately 60 measurement periods (with single transmission per 60-s period), with more distant nodes showing slightly reduced operational life due to increased communication load.
