## Supplementary Figure 6 for "Wireless Sensor Network: New Concept of Spatial-Temporal Monitoring Plant–Environment Interactions"

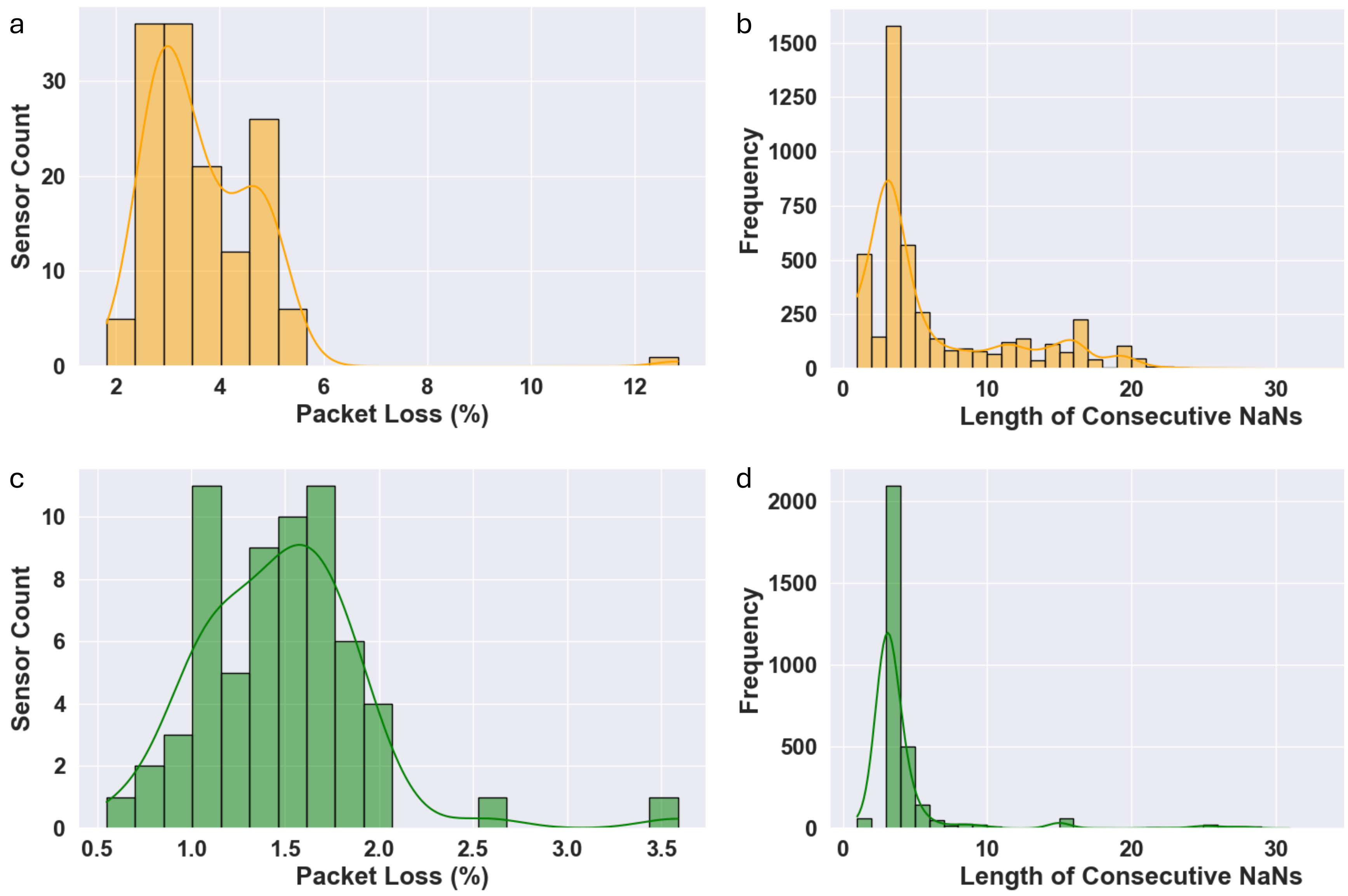


**Supplementary Figure 6. Packet loss and gap duration distributions in two greenhouse deployments. (**a–b) Polycarbonate greenhouse, 11 days, 144 sensor nodes ($7.55 \times 10^5$ expected 3-min records). (a) Histogram of packet loss per node (% of missing 3-min records), computed after de-duplication and filtering described in Methods. (b) Distribution of gap durations, where a gap is a consecutive run of missing 3-min records; the x-axis is the number of 3-min intervals per gap, the y-axis is the count of gaps.(c–d) Glasshouse, 30 days, 64 sensor nodes ($9.83 \times 10^5$ expected records). (c) Packet-loss histogram as in (a). (d) Gap-duration distribution as in (b). Both deployments used duplicate transmissions (two packets every 3 min); only the latest packet per interval was stored.
