## Supplementary Figure 7 for "Wireless Sensor Network: New Concept of Spatial-Temporal Monitoring Plant–Environment Interactions"

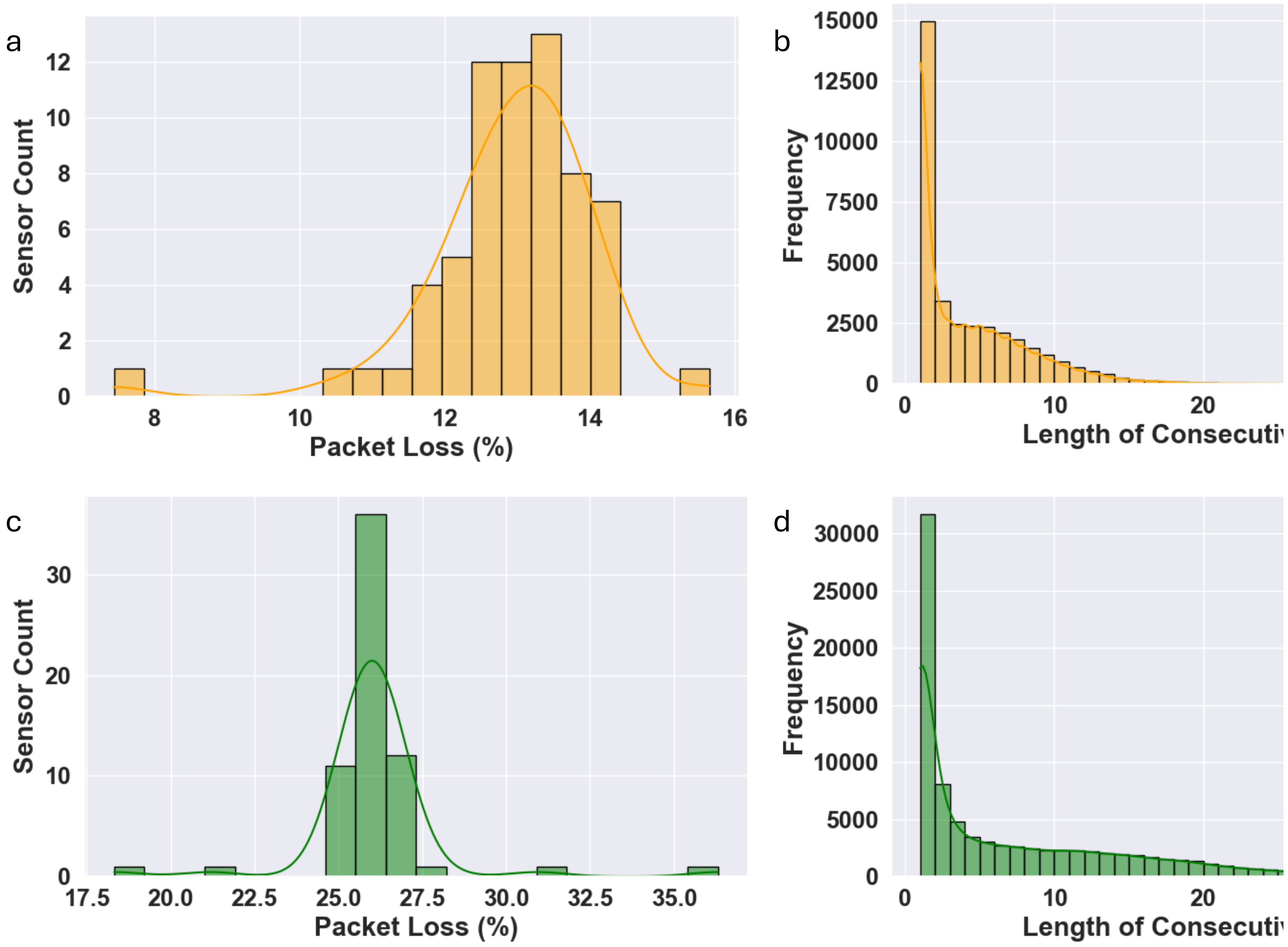


**Supplementary Figure 7. Packet loss and gap-duration distributions under single-transmission mode (one packet per 3 min). (a–b) 60-day growth-chamber deployment with 66 sensor nodes.** (a) Histogram of packet loss per node (% of missing 3-min records, see Methods). (b) Distribution of gap durations, where the x-axis is the number of consecutive missing 3-min records and the y-axis is the count of gaps. (c–d) 90-day deployment with 64 sensor nodes, same single-transmission configuration. (c) Packet-loss histogram as in (a). (d) Gap-duration distribution as in (b). Data processing matched the procedure described for Supplementary Fig. 6.
