## Supplementary Figure 8 for "Wireless Sensor Network: New Concept of Spatial-Temporal Monitoring Plant–Environment Interactions"

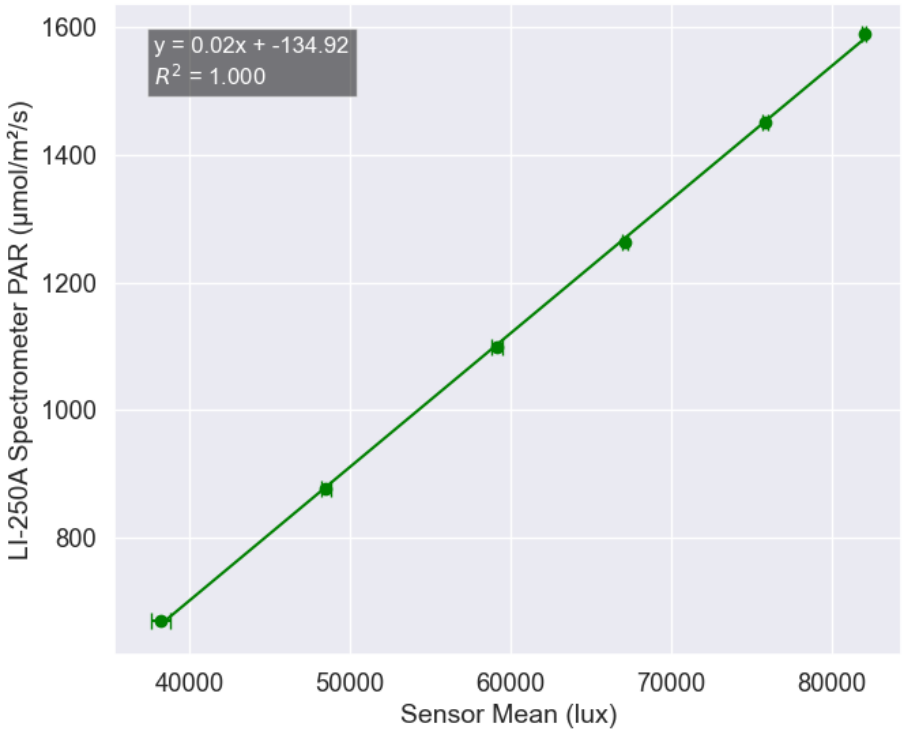


**Supplementary Figure 8. Correlation between SP light sensor readings and LI-COR LI-250A spectrometer measurements under growth-room lighting.**
Mean ± SE values from three SP sensors, compared with simultaneous measurements from a LI-COR LI-250A spectrometer across a light intensity range of 37,000–82,000 lux (corresponding to 670–1600 μmol $m^{-2}$ $s^{-1}$ PAR). The growth-room LED fixtures served as the light source. A strong linear correlation was observed ($R^2$ = 1.000). Unlike the calibration curve presented in Supplementary Figure 1D (derived under the Licor's 6400 red-blue LED spectrum), this calibration was performed under broad white-spectrum growth-room lighting, confirming that SP lux measurements are linearly related to PAR values across different spectral conditions. Error bars represent ±SE; where not visible, SE is smaller than the symbol size.
